# Nuclear adenosine metabolism defines a metabolic vulnerability unmasked by TP53 loss

**DOI:** 10.64898/2026.07.06.736733

**Authors:** Julia Urgel-Solas, Anna Benito-Ferrer, Laura Ortet, Savvas Kourtis, Etna Abad, Albert Coll Manzano, Anna Shevzov-Zebrun, Camilla Reiter Elbæk, Carlos M. Martínez, Matthew G Vander Heiden, Sara Sdelci, Ana Janic

**Affiliations:** Department of Medicine and Life Sciences, Pompeu Fabra University, Barcelona, Spain; Center for Genomic Regulation, Barcelona, Spain; Universitat de Barcelona, Barcelona, Spain; Instituto Murciano de Investigación Biosanitaria (IMIB-Pascual Parrilla), Murcia, Spain; Department of Biology, Massachusetts Institute of Technology, Cambridge, MA, USA; Koch Institute for Integrative Cancer Research, Massachusetts Institute of Technology, Cambridge, MA, USA; Broad Institute of MIT and Harvard, Cambridge, MA, USA; Dana-Farber Cancer Institute, Boston, MA, USA

**Author notes:** Equal contribution (Julia Urgel-Solas, Anna Benito-Ferrer).

## Abstract

Metabolic adaptation is essential for cells experiencing chronic genomic stress, yet how such adaptations are organized at the nuclear level remains poorly understood. Loss of TP53 is associated with elevated genomic instability, DNA damage and altered metabolic requirements, creating context specific dependencies. Here, we identify a requirement for *de novo* purine biosynthesis in TP53-deficient cells, with a pronounced dependence on adenosine related metabolism. Perturbation of purine synthesis increases DNA damage and reduces nuclear ATP availability, particularly in TP53-deficient cells, and is accompanied by a rapid increase in histone methylation. This chromatin response is also induced by acute DNA damage and occurs with fast kinetics, indicating that histone methylation is an early, intrinsic feature of the nuclear stress response rather than a secondary epigenetic remodeling. Interfering with histone methylation is associated with reduced nuclear ATP levels and impaired DNA damage resolution, linking chromatin state to nuclear energy homeostasis. Genetic disruption of the purine biosynthesis enzyme PFAS selectively impairs the growth of TP53-deficient tumours *in vivo*, establishing the physiological relevance of this metabolic dependency. Together, these findings reveal nuclear adenosine metabolism as a compartmentalised adaptive response to genotoxic stress and highlight chromatin associated methylation as a key feature of nuclear metabolic regulation unmasked by TP53 loss.

## Main

Maintenance of genome integrity imposes substantial energetic and metabolic demands on cells, particularly during DNA replication and repair^1^. Genetic lesions that compromise genome surveillance frequently drive compensatory metabolic adaptations, creating context specific dependencies that shape cell fitness under stress^2^. Understanding how these dependencies arise is central to both cancer biology and fundamental principles of metabolic regulation.

Among the most frequently altered regulators of genome stability is the tumour suppressor TP53^3,4^. Beyond its canonical roles in cell cycle arrest, apoptosis, and senescence^5,6^, TP53 coordinates metabolic programs that align energy availability with genome associated processes. Wild-type TP53 promotes oxidative metabolism, restrains glycolysis, and modulates signalling pathways such as mTOR and AMPK to balance biosynthesis and energy production^7–9^. Loss of TP53 disrupts this coordination and is associated with elevated basal DNA damage, replication stress, and persistent activation of genome maintenance pathways^10^. Accordingly, TP53-deficient cells undergo extensive metabolic reprogramming that supports proliferation despite genomic instability and oncogenic stress^11^.

To date, metabolic adaptations associated with TP53 loss have been characterised primarily at the level of cytoplasmic and mitochondrial pathways, including glucose metabolism, oxidative phosphorylation, and anabolic biosynthesis^12–14^. Which metabolic pathways become selectively required to cope with the increased energetic burden imposed by TP53 deficiency, and how these requirements relate to genome maintenance, remain incompletely understood.

Here, we set out to identify metabolic vulnerabilities associated with TP53 loss. Using a CRISPR-Cas9 functional genetic screen, we uncover a selective dependency on *de novo* purine biosynthesis, with a pronounced requirement for adenosine related metabolism. This dependency is mediated by core components of the purine synthesis pathway, including PFAS, whose genetic depletion selectively impairs the growth of TP53-deficient tumours *in vivo*. Pharmacological inhibition of purine synthesis similarly exposes a vulnerability that is rescued by adenosine but not by guanosine, indicating a specific reliance on the adenosine branch of purine metabolism.

Mechanistic interrogation of this dependency revealed an unexpected link to nuclear metabolism. Nuclear processes such as DNA repair, transcription, and chromatin remodelling impose high and dynamic energetic demands, yet have generally been assumed to rely on metabolites supplied from the cytoplasm, with limited consideration of compartment specific regulation.

We find that inhibition of *de novo* purine synthesis in TP53-deficient cells that are dependent on adenosine, is associated with increased DNA damage, induction of histone methylation, and reduced nuclear ATP availability. To distinguish whether histone methylation reflects a secondary epigenetic adaptation or an intrinsic DNA damage response, we examined its kinetics following acute genotoxic stress. Low dose γ-irradiation induces a peak of DNA damage within hours, and within this short time frame histone methylation is similarly increased, indicating that methylation is engaged rapidly in response to DNA damage rather than arising from slower epigenetic remodelling processes. Interfering with histone methylation under these conditions is associated with reduced nuclear ATP levels and delayed DNA damage resolution. Together, these observations support an intimate connection between chromatin modification and nuclear energy homeostasis and are consistent with the idea that chromatin associated processes may participate in adenosine recycling when *de novo* synthesis is limited.

Collectively, our findings identify nuclear adenosine metabolism as a critical adaptive response unmasked by TP53 loss. By linking purine synthesis dependency, chromatin modification, and nuclear energy balance, this work reveals a previously unrecognised mode of compartmentalised metabolic regulation that supports cell fitness under genotoxic stress. Given the prevalence of TP53 alterations across human cancers, these findings have broader implications for understanding how metabolic vulnerabilities arise from chronic genome instability and how nuclear metabolism contributes to tumour biology.

## Results

### Loss of TP53 creates a selective dependency on purine synthesis

TP53 and KRAS are among the most frequently mutated genes in non–small-cell lung cancer^15–17^. Across multiple cancer types, patients harbouring concurrent KRAS mutations and TP53 loss display significantly poorer overall survival compared to patients carrying only one of these alterations or neither (Figure 1A and Supplementary Data 1). This genetic context is associated with elevated genomic instability and increased cellular stress^18,19^, providing a suitable framework to interrogate how metabolic pathways support cellular fitness under such conditions. To identify metabolic dependencies specifically associated with TP53 loss in a KRAS-mutant background, we performed a focused CRISPR-Cas9 genetic screen targeting metabolic pathways in isogenic lung cancer cell models. We employed a human metabolism-focused CRISPR knockout library comprising approximately 30,000 sgRNAs targeting around 3,000 metabolic enzymes, small molecule transporters, and metabolism-associated transcriptional regulators, together with control sgRNAs, delivered in a Cas9-expressing lentiviral vector^20^. To ensure that identified dependencies were attributable to TP53 status rather than differences in oncogenic signalling, we used an isogenic system based on human A549 lung adenocarcinoma TP53 wild type cells harbouring an endogenous KRAS^G12S^ mutation (K^A549^) and an A549-derived counterpart with TP53 deletion (KP^A549^)^21^ (Figure 1B). Importantly, K^A549^ and KP^A549^ cells proliferated at comparable rates under standard culture conditions (Supplementary Figure 1A), supporting their use as a matched system for identifying genotype dependent metabolic dependencies. K^A549^ and KP^A549^ cells were transduced with the metabolic sgRNA library, selected with antibiotic for six days, and cultured for an additional fourteen days prior to genomic DNA extraction and sgRNA quantification (Figure 1B). Quality control analyses confirmed robust screen performance, including consistent Gini indices, minimal sgRNA dropout, uniform sgRNA representation, and clear separation of samples by condition in principal component analysis (PCA) (Supplementary Figure 1B-E and Supplementary Data 2). As an internal validation, sgRNAs targeting the tumour suppressor PTEN were selectively enriched in TP53-deficient cells, consistent with previous reports that combined loss of TP53 and PTEN confers a proliferative advantage (Figure 1C)^22^. Across both cell lines, we identified 69 genes whose sgRNAs were significantly depleted, indicating a requirement for proliferation independent of TP53 status (Figure 1C and Supplementary Data 3). These genes were enriched for pathways associated with mitochondrial function, energy production, vesicle organisation, proton transport, and autophagy, reflecting core cellular processes broadly required for viability (Supplementary Figure 1F). In contrast, a distinct set of approximately 65 genes exhibited selective sgRNA depletion exclusively in TP53-deficient cells (average relative depletion score < -0.45), indicating a dependency that emerges specifically upon TP53 loss (Supplementary Data 3). Gene Ontology analysis of this TP53-dependent gene set revealed a strong and selective enrichment for the *de novo* purine biosynthesis pathway (Figure 1C-D and Supplementary Figure 1G). Notably, sgRNAs targeting multiple enzymes across the pathway were depleted, including phosphoribosyl pyrophosphate amidotransferase (PPAT), phosphoribosylaminoimidazole carboxylase and phosphoribosylaminoimidazolesuccinocarboxamide synthetase (PAICS), phosphoribosylglycinamide formyltransferase, phosphoribosylglycinamide synthetase, phosphoribosylaminoimidazole synthetase (GART), adenylosuccinate lyase (ADSL), and phosphoribosylformylglycinamidine synthase (PFAS) (Figure 1E-F). In contrast, enzymes involved in *de novo* pyrimidine biosynthesis were not overrepresented among TP53-specific dependencies (Figure 1C-F and Supplementary Figure 1G), indicating that the observed phenotype is not a general consequence of increased nucleotide demand associated with proliferation. To assess whether this dependency extends beyond the isogenic A549 system, we analysed genome-wide essentiality data from the DepMap project^23^ and integrated these with transcriptomic data from the Cancer Cell Line Encyclopedia (CCLE)^24^. Cell lines were stratified according to TP53 activity using the PROGENy algorithm^25^, which infers pathway activity based on downstream transcriptional signatures. Across diverse cancer cell lines, enzymes involved in de novo purine biosynthesis exhibited increased essentiality in cells with low inferred TP53 activity, resembling TP53-deficient states (Figure 1G and Supplementary Data 4). Consistent with this, analysis of The Cancer Genome Atlas (TCGA) Pan-Cancer datasets^26^ revealed higher expression of purine biosynthesis enzymes, including PPAT, GART, PFAS, and PAICS, in TP53-mutant tumours compared to TP53-proficient tumours across multiple cancer types (Figure 1H).

**Figure 1.**
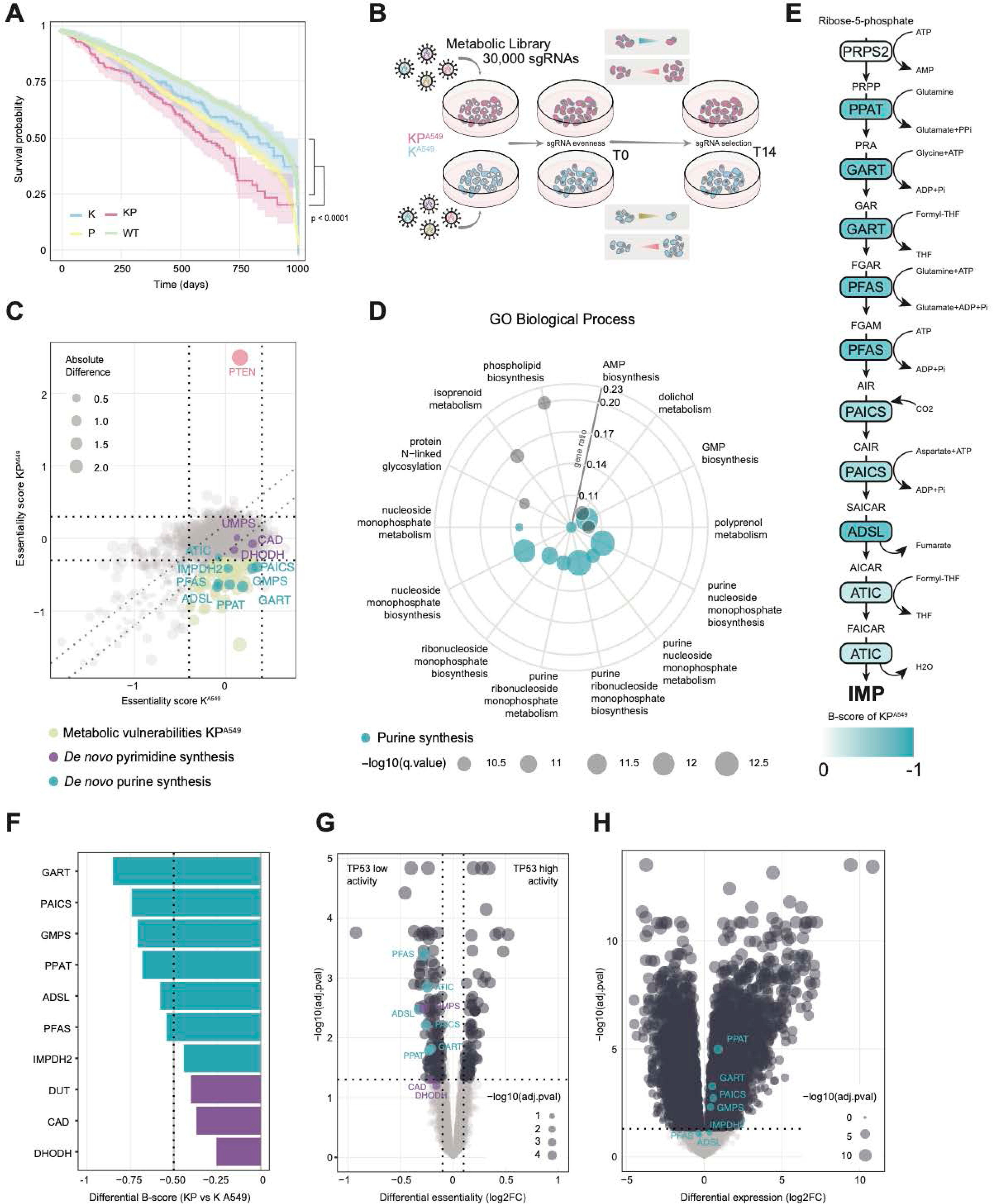
Purine metabolism is essential in TP53-deficient cells. A) Kaplan–Meier curves showing patient overall survival. Data were retrieved from the TCGA-PANCAN dataset, with patients stratified based on KRAS and TP53 status. B) Schematic representation of the drop-out screen. C) Differentially abundant sgRNAs compared to day 0. Each dot represents the mean of all sgRNAs/gene. ß-scores (essentiality scores) are negative when a gene is more essential. In yellow, all the genes scored more essential in KP^A549^ cells than K^A549^ cells. In cyan, top scored purine synthesis genes in KP^A549^; in purple, pyrimidine metabolism-related genes. D) GO terms associated with significantly depleted sgRNAs in KP^A549^ vs K^A549^ lung cancer cells. E) KP^A549^ top-scored purine biosynthesis genes with their corresponding ß-scores. F) Differential essentiality scores from K^A549^ vs KP^A549^. In cyan de novo purine synthesis enzymes, in purple de novo pyrimidine synthesis enzymes. G) Project Achilles cancer cell lines (CCLE dataset) segregated by TP53 activity (Progeny). In cyan top-scored purine synthesis genes. P-value was calculated by two-sided, unpaired Student’s t-test performed for each gene individually and corrected for multiple comparisons using Benjamini-Hochberg (FDR) adjustment. H) Volcano plot showing differentially expressed genes between TP53-mutant and TP53-wild-type groups based on TCGA transcriptomics dataset (only stage III KRAS mutated tumours). Genes were considered significant based on a Benjamini-Hochberg adjusted p-value (FDR < 0.05) and log fold-change thresholds. Significantly upregulated purine biosynthesis genes in patients harbouring concurrent mutations in TP53 and KRAS are highlighted.

Together, these data identify de novo purine biosynthesis as a metabolic pathway that becomes selectively required upon TP53 loss in KRAS-driven lung cancer cells, suggesting that TP53 deficiency exposes a context dependent reliance on purine metabolism that supports cellular fitness under conditions of elevated genomic stress.

### TP53 loss drives sensitivity to purine synthesis inhibition that is selectively rescued by adenosine

Given the strong enrichment of purine metabolism genes identified in our genetic screen, we next examined the functional contribution of this pathway to cellular fitness in lung cancer models differing in TP53 status. In addition to the A549 isogenic pair (K^A549^ and KP^A549^), we used HBEC-3KT-derived human lung cancer cell lines (3KT) harbouring an oncogenic KRAS^G12D^ mutation that either retain TP53 (K^3KT^) or lack TP53 (KP^3KT^)^21^. These complementary systems allowed us to assess whether dependence on purine synthesis is a general feature associated with TP53 loss rather than a cell line specific effect.

To test requirement on *de novo* purine synthesis, we treated A549 and 3KT TP53-proficient and TP53-deficient cells with the purine synthesis inhibitor 6-mercaptopurine (6MP) (Figure 2A). Cell viability was quantified by automated nuclear counting using high-throughput immunofluorescence (HT-IF), as cells constitutively expressed a GFP nuclear localization signal (NLS) reporter. Across both cellular lung cancer models, TP53-deficient cells exhibited significantly increased sensitivity to 6MP compared to their TP53-proficient counterparts (Figure 2B,C and Supplementary Data 5), indicating that loss of TP53 enhances dependence on purine synthesis for proliferation.

**Figure 2.**
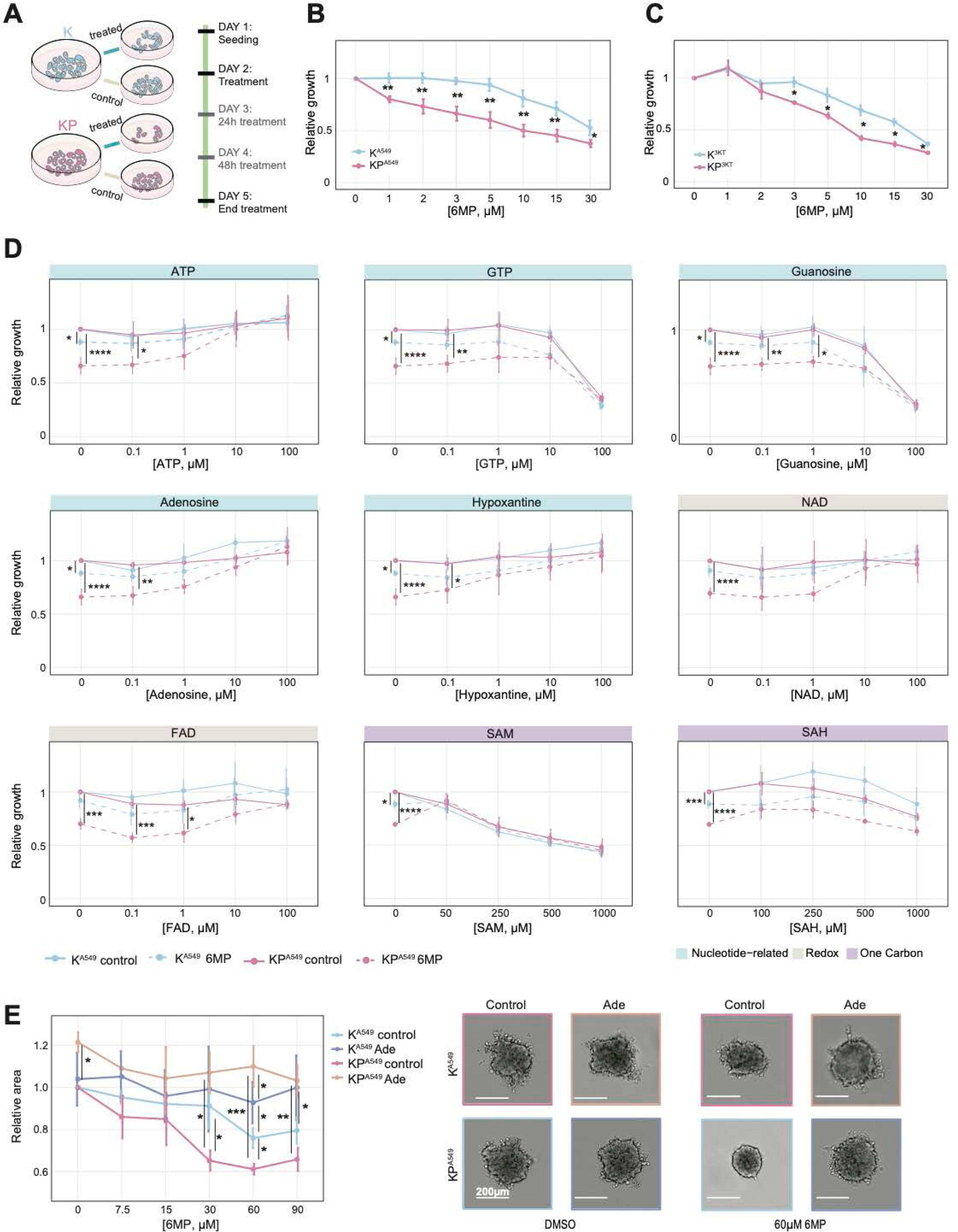
TP53-deficient cells are more sensitive to inhibition of purine synthesis. A) Schematic representation of the treatment administration regime. Relative growth of B) K^A549^ and KP^A549^, C) K^3KT^ and KP^KT^ cells treated with the indicated concentrations of 6MP or DMSO, and D) K^A549^ and KP^A549^ cells treated with DMSO or 6MP and supplemented with the indicated metabolites for 72 hours, following NLS-GFP signal analysis of nuclei count. E) (left) Relative spheroid area growth of K^A549^ and KP^A549^ treated with DMSO or 6MP for 7 days with the indicated concentrations, in presence or absence of 100µM of adenosine. (right) Representative images of spheroids, scale bar 200 μm. Data are normalised to the DMSO-treated condition. Data represents N=3 independent experiments. Mean ± SD, P-values ****P ≤ 0.00005; ***P ≤ 0.0005; **P ≤ 0.005; *P ≤ 0.05, pairwise t-test with Bonferroni adjusted.

To assess whether this sensitivity extends beyond lung cancer models, we examined murine pancreatic ductal adenocarcinoma cells derived from genetically engineered mouse models expressing oncogenic Kras^G12D^ with either wild-type Trp53 (KC^PDAC^), Trp53 deletion (KPC^PDAC^), or Trp53 missense mutations (KPC^R270H^ and KPC^R172H^)^27,28^. Consistent with the lung cancer data, Trp53-null or mutant PDAC cells displayed increased sensitivity to 6MP relative to Trp53-proficient cells (Supplementary Figure 2A-B and Supplementary Data 6), suggesting that TP53 loss broadly enhances sensitivity to purine synthesis inhibition across distinct epithelial cancer contexts.

Because 6MP broadly inhibits de novo purine synthesis, we next sought to distinguish whether the observed phenotype reflected a dependency on guanosine or adenosine metabolism. To selectively impair guanosine synthesis, we inhibited inosine monophosphate dehydrogenase using mycophenolic acid (MPA). In contrast to 6MP, MPA treatment did not preferentially reduce viability in TP53-deficient cells. Notably, in the A549 isogenic system, TP53-deficient cells displayed increased resistance to MPA compared to TP53-proficient cells (Supplementary Figure 2C and Supplementary Data 7). These results indicate that impaired guanosine synthesis alone is insufficient to account for the sensitivity of TP53-deficient cells to purine synthesis inhibition and instead point toward a selective reliance on adenosine metabolism.

To directly test this possibility, we examined whether supplementation with purine metabolites could rescue proliferation under 6MP treatment. Supplementation with ATP restored proliferation of TP53-deficient KP^A549^ cells to levels comparable to TP53-proficient cells (Figure 2D and Supplementary Data 8). In contrast, GTP supplementation failed to rescue proliferation and was toxic at higher concentrations (Figure 2D). Because extracellular nucleotides are primarily taken up as nucleosides, we next supplemented cultures with adenosine or guanosine. Adenosine supplementation fully recapitulated the rescue observed with ATP, whereas guanosine failed to restore proliferation (Figure 2D and Supplementary Data 8).

Consistent with this specificity, supplementation with additional adenosine related metabolites, including hypoxanthine, nicotinamide adenine dinucleotide, and flavin adenine dinucleotide, also rescued proliferation under 6MP treatment (Figure 2D and Supplementary Data 8). In contrast, S-adenosyl-L-methionine (SAM) and S-adenosyl-L-homocysteine (SAH) failed to restore viability despite containing an adenosine moiety. Similar results were observed in spheroid cell cultures, where adenosine supplementation restored the growth of TP53-deficient spheroids treated with 6MP to levels comparable to TP53-proficient controls (Figure 2E and Supplementary Data 9). As expected, supplementation with pyrimidine nucleotides did not rescue growth inhibition induced by 6MP (Supplementary Figure 2D and Supplementary Data 10).

Together, these results indicate that TP53 loss confers a selective dependence on *de novo* purine synthesis that is specifically linked to adenosine metabolism, rather than to guanosine or pyrimidine nucleotide production.

### TP53 deficiency alters purine metabolic homeostasis

To investigate the metabolic basis underlying increased reliance on adenosine metabolism, we performed quantitative targeted metabolomic profiling in TP53-proficient and TP53-deficient 3KT cells. Using mass spectrometry, we quantified 111 metabolites spanning ten metabolic pathways (Supplementary Data 11). PCA revealed clear separation between TP53-proficient and TP53-deficient cells, indicating distinct metabolic states (Supplementary Figure 3A). Direct comparison of metabolite abundances identified widespread metabolic differences (Supplementary Figure 3B and Supplementary Data 12), with the most prominent changes observed within nucleoside- and nucleotide-associated pathways (Figure 3A-B).

**Figure 3.**
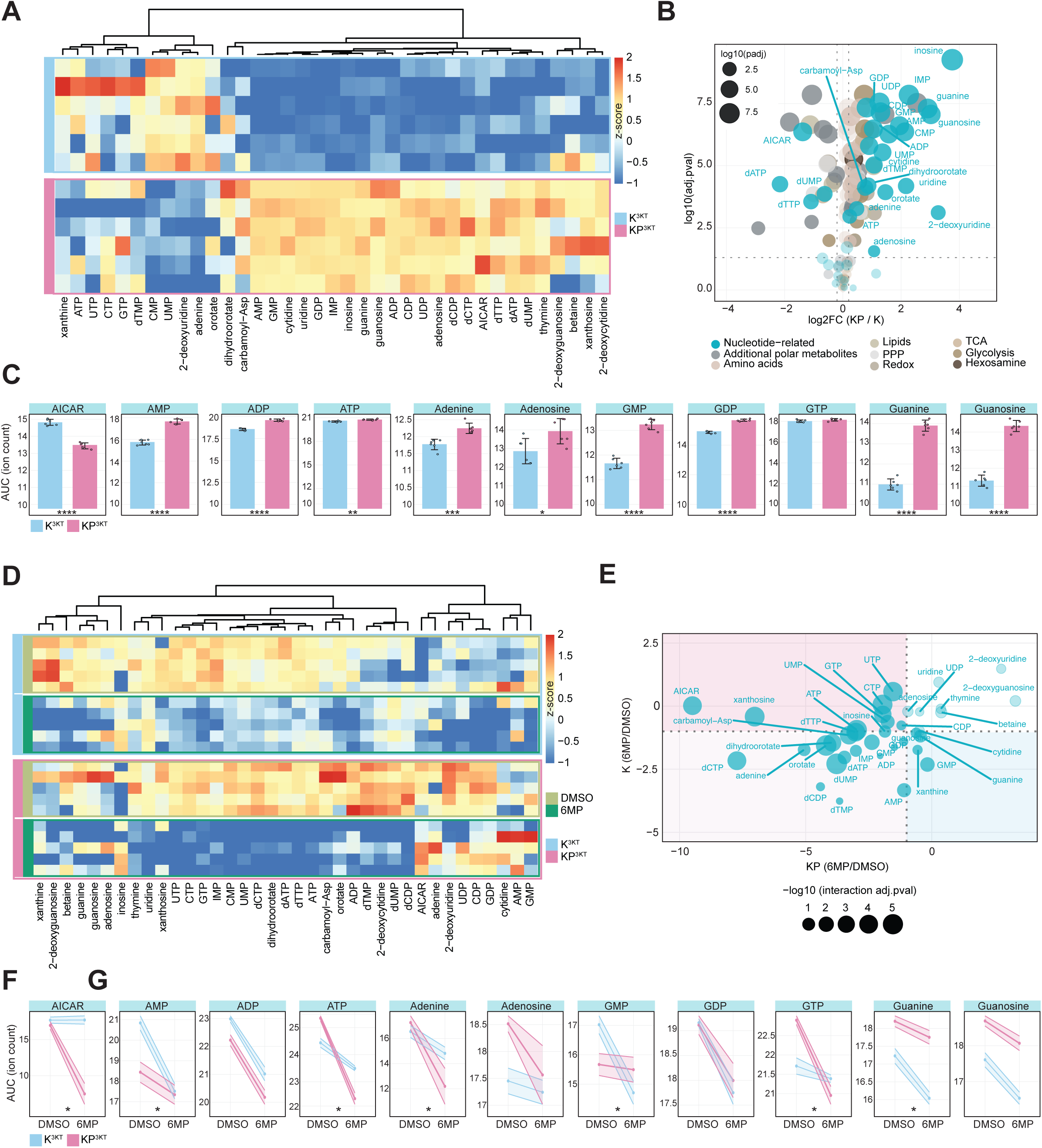
TP53 loss alters *de novo* purine metabolome. A) Hierarchical clustering heatmap of nucleotide and related metabolite Z-scores in K^3KT^ and KP^3KT^ cells. B) Volcano plot showing normalised differential (K^3KT^ vs KP^3KT^ cells) abundance of metabolites. Cyan circles indicate nucleotide-related metabolites, while circle size reflects –log10(p-adjusted). C) Corrected area under the curve (AUC) of ion counts for purine nucleotides (AICAR, AMP, ADP, ATP, adenine, adenosine, GMP, GDP, GTP, guanine, guanosine). D) Heatmap of metabolite Z-scores in K^3KT^ and KP^3KT^ cells treated with DMSO or 6MP 5 µM for 72 hours. E) Interaction volcano plot displaying differential response to 6MP between K^3KT^ and KP^3KT^cells; dot size reflects statistical significance of interaction. F,G) Relative quantification of purine metabolites after 72 hours treatment with DMSO or 5 µM 6MP in K^3KT^ and KP^3KT^ cells. Data represent N=5 (baseline condition A-C) or N=6 (treatment condition D-G) independent experiments. Mean ± SD. P-values: ****P ≤ 0.00005; ***P ≤ 0.0005; **P ≤ 0.005; *P ≤ 0.05, pairwise t-test with Bonferroni adjustment.

Notably, TP53-deficient cells exhibited increased levels of purine nucleotides, accompanied by a marked reduction in 5-aminoimidazole-4-carboxamide ribonucleotide (AICAR), a key intermediate in *de novo* purine biosynthesis (Figure 3C). This pattern is consistent with enhanced use of AICAR to sustain purine production. Guanine, guanosine, and selected pyrimidine metabolites were modestly elevated (Figure 3A-B and Supplementary Figure 3B). No significant differences in cell cycle distribution were observed between TP53-proficient and TP53-deficient cells (Supplementary Figure 3C), indicating that these metabolic adaptations are sufficient to support cell cycle progression under basal conditions.

To assess how these cells respond to purine deprivation, we performed metabolomic profiling following 72 hours of 6MP treatment (Supplementary Data 13). PCA revealed distinct clustering of treated and untreated samples in both cell lines, with TP53-deficient cells exhibiting a more pronounced metabolic shift in response to 6MP (Supplementary Figure 3D). As expected, 6MP treatment led to a strong reduction in purine nucleotide levels in both TP53-proficient and TP53-deficient cells (Figure 3D). Pyrimidine metabolites were also reduced, potentially reflecting compensatory mechanisms aimed at maintaining nucleotide balance (Figure 3D-E, Supplementary Figure 3E and Supplementary Data 14).

Strikingly, AICAR levels were profoundly depleted in TP53-deficient cells following 6MP treatment, whereas AICAR remained relatively stable in TP53-proficient cells (Figure 3E-F). This collapse of AICAR was accompanied by a reduction in downstream purine metabolites (Figure 3G). Given the increased dependence of TP53-deficient cells on purine biosynthesis genes, this exaggerated depletion suggests an impaired ability to maintain purine metabolic homeostasis under *de novo* synthesis inhibition, providing an explanation for their increased sensitivity to 6MP compared to TP53 wild-type derivatives.

### Adenosine dependency driven by TP53 loss links DNA damage to histone hypermethylation

The depletion of TP53, widely regarded as the guardian of the genome, compromises the DNA damage response and genome integrity^29^. We therefore hypothesised that the increased dependence of TP53-deficient cells on adenosine related biosynthetic pathways reflects an enhanced requirement to cope with chronic DNA damage arising from TP53 loss. As perturbations in nucleotide availability can directly affect DNA replication and repair, we further reasoned that altered purine metabolism in TP53-deficient cells might exacerbate replication stress, thereby amplifying DNA damage and downstream chromatin alterations.

To test this hypothesis, we quantified DNA damage in the A549 isogenic pair under basal conditions and following inhibition of *de novo* purine synthesis with 6MP and assessed the effect of adenosine supplementation. Given the strong influence of cell-cycle progression on DNA damage accumulation, we analysed the data in a cell-cycle resolved manner, assigning individual cells to specific cell-cycle phases based on nuclear area and DAPI intensity, as previously described^30–32^. DNA damage was assessed by HT-IF detection of γH2AX and 53BP1 foci, two established markers of DNA double-strand breaks^33,34^ (Supplementary Data 15). Under basal conditions and upon treatment with 6MP, TP53-deficient 3KT and A549 lung cancer cells exhibited higher levels of DNA damage, as demonstrated by increased γH2AX and 53BP1 accumulation, compared to their TP53-proficient counterparts, with DNA damage further intensified with cell-cycle progression. Notably, adenosine supplementation significantly reduced 6MP-induced DNA damage in both TP53-deficient and TP53-proficient lung cancer cells, and in most conditions lowered γH2AX and 53BP1 levels near to baseline (Figure 4A-B and Supplementary Figure 4 A-B; Supplementary Data 16).

**Figure 4.**
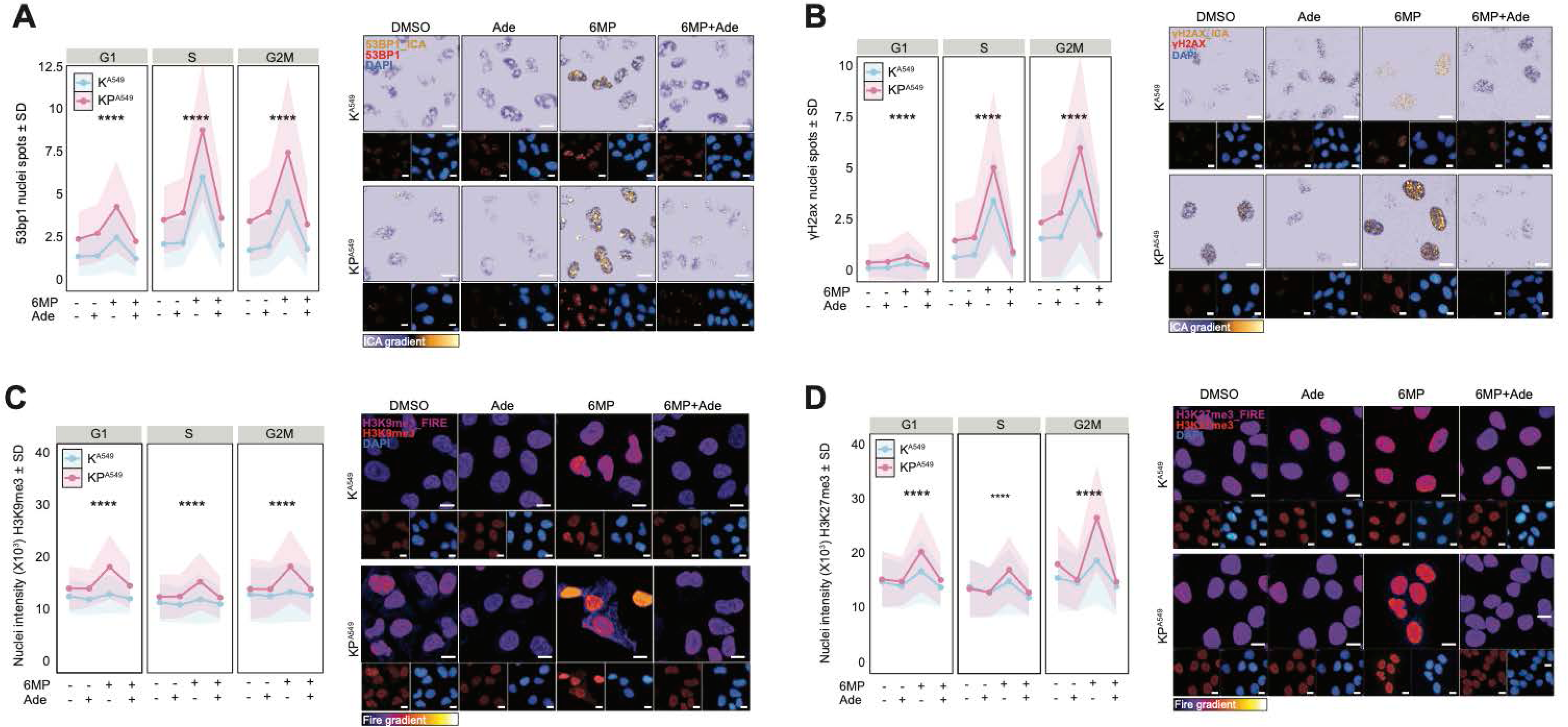
Purine depletion induces DNA damage and histone methylation preferentially in TP53-deficient lung cancer cells. K^A549^ and KP^A549^ treated with DMSO or 15 µM 6MP for 72 hours in presence or absence of 100 µM Adenosine. A) (left) Quantification of 53BP1 foci, (right) representative images. B) (left) Quantification of γH2AX foci, (right) representative images. N = 1000-4000 cells per condition. Mean ± SD, Bonferroni adjusted one-way ANOVA, ****P ≤ 0.00005. 53BP1 and γH2AX are in ICA_gradient and DAPI marks nuclei in blue. Scale bar = 10 µm. C) (left) Quantification of H3K9me3 nuclear intensity in K^A549^ and KP^A549^, (right) representative images, D) (left) Quantification of H3K27me3 nuclear intensity in K^A549^ and KP^A549^, (right) representative images. N = 1000-4000 cells per condition. Data represents N=2 independent experiments. Mean ± SD, Bonferroni adjusted one-way ANOVA, ****P ≤ 0.00005. H3K9me3 and H3K27me3 are in FIRE_gradient, DAPI marks nuclei in blue; scale bar = 10 µm.

Consistent with the observation that the strongest differences between TP53-proficient and TP53-deficient cells arise from S phase onwards, we directly examined DNA replication dynamics using DNA fibre assays in K^3KT^ and KP^3KT^ cells treated with 6MP (Supplementary Figure 4C and Supplementary Data 17). Under basal conditions, KP^3KT^ cells displayed significantly higher replication fork speed than K^3KT^ cells, as indicated by longer DNA fibres, revealing accelerated replication dynamics in the absence of TP53. 6MP treatment caused a decrease in DNA replication speed in both, K^3KT^ and KP^3KT^ cells, with a significantly more pronounced drop in KP^3KT^. Adenosine supplementation partially rescued 6MP induced replication fork speed impairment in both genetic backgrounds (Supplementary Figure 4C). Given that imbalances in nucleotide availability are a well established source of replication stress^35^, these data indicate that TP53-deficient cells, which replicate faster at baseline, are particularly vulnerable to purine limitation during DNA synthesis.

Because histone methylation is a chromatin modification that is directly linked to cellular metabolic state^36^ and is engaged during DNA damage responses^37^, we examined whether purine metabolism inhibition was accompanied by changes in histone methylation in TP53-deficient cells. We measured H3K9me3 and H3K27me3 levels, markers of constitutive and facultative heterochromatin^38^, in TP53-proficient and -deficient A549 cells treated with 6MP, with or without adenosine supplementation (Supplementary Data 18). While basal levels of both marks were comparable between untreated K^A549^ and KP^A549^ cells, 6MP treatment induced a robust increase in H3K9me3 and H3K27me3 specifically in TP53-deficient A549 cells across all cell-cycle phases. This hypermethylation phenotype was effectively rescued by adenosine supplementation (Figure 4C–D). Similar results were observed in the K^3KT^/KP^3KT^ isogenic system (Supplementary Figure 4D–E; Supplementary Data 19) and were not explained by increased expression of canonical histone methyltransferases in TP53-deficient cells (Supplementary Figure 4F and Supplementary Data 20).

Together, these data indicate that TP53-deficient cells under baseline conditions have elevated replication stress and DNA damage, as previously shown, which renders them vulnerable to disruptions in purine metabolism. Inhibition of *de novo* purine synthesis further exacerbates replication stress, leading to excessive DNA damage and a concomitant increase in repressive histone methylation. Thus, the sensitivity of TP53-deficient cells to purine synthesis inhibition reflects a failure to buffer replication associated DNA damage, resulting in both genomic and epigenetic dysregulation.

### TP53 loss induces DNA methylation to support nuclear adenosine recycling

Given the combined effects of purine synthesis inhibition on DNA damage, nuclear ATP levels, and histone methylation in TP53-deficient cells, we next examined whether these nuclear responses were mechanistically linked. We hypothesised that TP53 loss creates a compartment specific vulnerability in nuclear ATP availability, particularly under conditions of nucleotide stress. This vulnerability likely arises from the combined burden of accelerated DNA replication and chronically elevated DNA damage, both of which impose high local ATP demand within the nucleus to sustain genome duplication and repair^39^.

To test this, we transduced K^A549^ and KP^A549^ cells with genetically encoded ATP sensors targeted to either the cytoplasm or the nucleus^40,41^ and measured ATP levels following inhibition of *de novo* purine synthesis with 6MP (Supplementary Data 21). Cytoplasmic ATP levels were largely unaffected by 6MP in both genotypes (Figure 5A). In contrast, nuclear ATP levels were selectively and significantly reduced in TP53-deficient A549 cells, while remaining stable in TP53-wild type A549 cells (Figure 5B). These results indicate that TP53 loss is associated with an impaired ability to sustain nuclear ATP pools under purine synthesis inhibition. We reasoned that reduced nuclear ATP availability would necessitate activation of local adenosine recycling pathways within the nucleus. A major source of nuclear adenosine can come from DNA and histone methylation^42^, during which SAM, the universal methyl donor^43^, is converted into SAH, subsequently hydrolysed by SAH hydrolase (SAHH, e.g. AHCY) to generate homocysteine and adenosine^44^. Metabolomic profiling revealed that SAM levels were comparable between KP^3KT^ and K^3KT^ isogenic lung cell lines and decreased similarly following 6MP treatment, whereas SAH levels were reduced to greater extent in KP^3KT^ cells compared to K^3KT^ cells (Figure 5C). This selective depletion of SAH is in consistent with the increased histone methylation observed in TP53-deficient cells and suggests a potential link between chromatin methylation and adenosine recycling within the nucleus.

**Figure 5.**
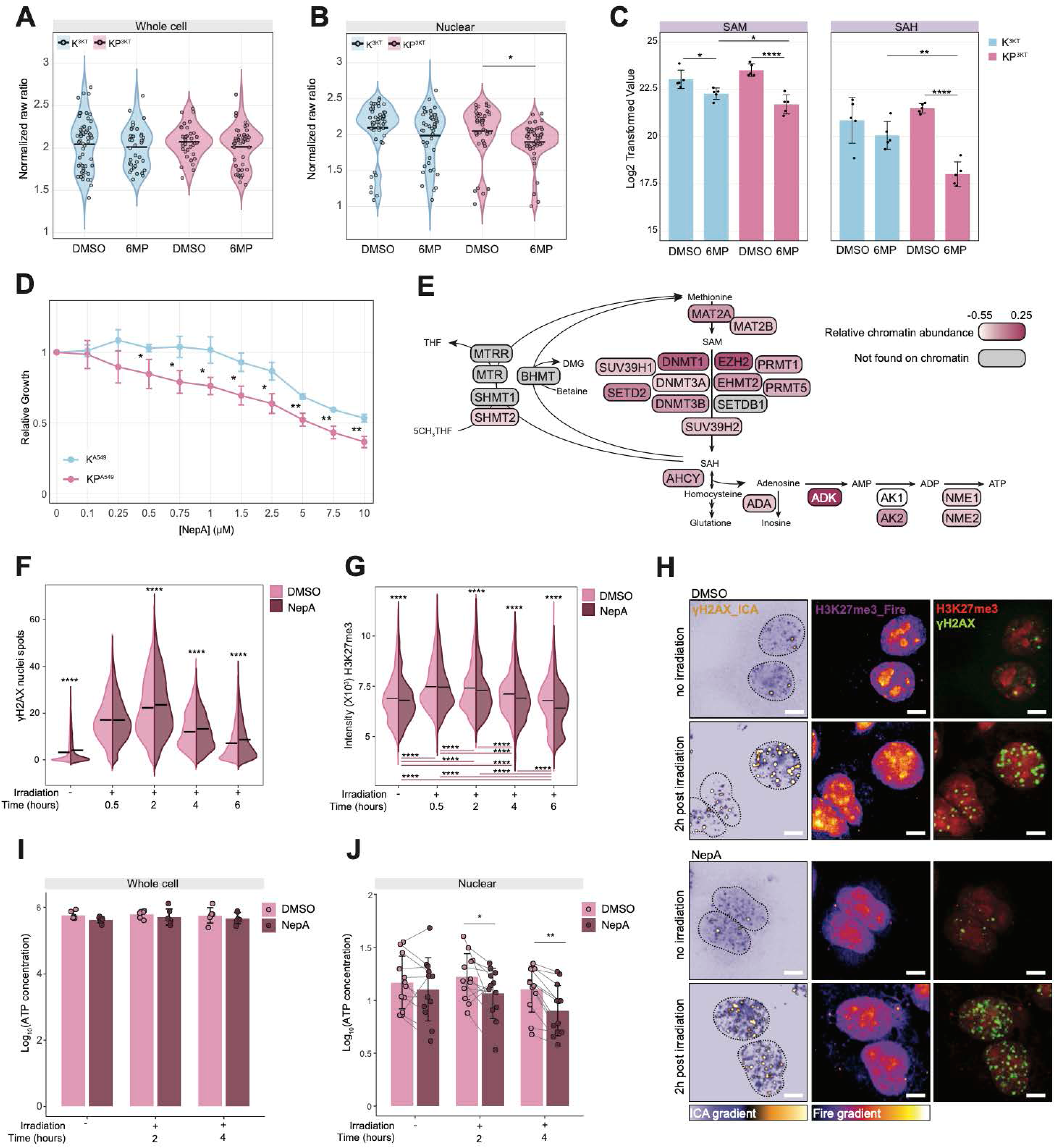
TP53 loss induces DNA methylation to recycle nuclear adenosine. A,B) ATP-sensor quantification in K^A549^ and KP^A549^, normalised to total expression of the vector for both cytoplasmic ATP (A) and nuclear ATP (B). C) Normalised relative abundance of SAM and SAH in K^3KT^ and KP^3KT^ cells with DMSO or 5 µM 6MP. D) Growth of K^A549^ and KP^A549^ treated with the indicated concentrations of NepA for 72 hours. Data is represented as mean ± SD, P-values ****P ≤ 0.00005; ***P ≤ 0.0005; **P ≤ 0.005; *P ≤ 0.05, pairwise t-test with Bonferroni adjusted. E) Scheme of the enzymes involved in SAM associated with adenosine and ATP metabolism, highlighting their relative chromatin abundance (pink gradient). F) Quantification of γH2AX foci of KP^A549^ cells treated with 10 µM NepA or DMSO and combined with or without γ-irradiation at the indicated timepoints. Two-way ANOVA with Holm-adjusted pairwise comparisons, ****P ≤ 0.00005. G) Quantification of high-intensity regions of H3K27me3 of KP^A549^ cells treated with 10 µM NepA or DMSO and combined with or without γ-irradiation at the indicated timepoints. Two-way ANOVA with Holm-adjusted pairwise comparisons, ****P ≤ 0.00005. H) Representative images of γH2AX in ICA_gradient and H3K27me3 in FIRE_gradient. Scale bar = 10 µm. I) Whole cell and J) nuclear fractions ATP measurements of KP^A549^ following DMSO or 10µM NepA treatment with or without irradiation. Grey lines show the replicates. Linear modeling with Batch as a covariate, followed by Holm-adjusted pairwise comparisons, **P ≤ 0.005; *P ≤ 0.05.

In line with this model, pharmacological repression of SAH hydrolysis using the AHCY inhibitor neplanocin A (NepA) produced a significantly stronger antiproliferative effect in KP^A549^ cells than in K^A549^ cells (Figure 5D; Supplementary Data 22), indicating a stronger reliance of TP53-deficient cells on methylation derived adenosine recycling. To support adenosine availability and ATP production in the nucleus, enzymes linking methylation reactions to adenosine and ATP metabolism would be expected to localise to the nuclear compartment. Using chromatome proteomics^45^, we found that multiple enzymes involved in SAM-related adenosine and ATP metabolism are physically associated with chromatin and selectively enriched in TP53-mutant cells (Figure 5E and Supplementary Figure 5A). These include SUV39H1 and EZH2, the methyltransferases responsible for depositing the H3K9me3 and H3K27me3 histone marks that increase upon 6MP treatment, as well as AHCY, adenosine kinase (ADK), and adenylate kinase 2 (AK2) (Supplementary Figure 5B). Together, the chromatin association of these enzymes supports the existence of a spatially confined nuclear program linking histone methylation to adenosine and ATP metabolism. To assess how DNA damage influences histone methylation and nuclear ATP availability independently of prolonged purine synthesis inhibition, TP53-deficient A549 cells were exposed to a single low-dose γ-irradiation (2 Gy) with or without NepA. This dose induces transient and fully repairable DNA damage (Supplementary Figure 5C), enabling dynamic assessment of early DNA damage repair. Cells were analysed at 0.5, 2, 4, and 6 hours post-irradiation to quantify DNA damage accumulation (γH2AX and 53BP1) and induction of the repressive histone methylation mark H3K27me3. Following irradiation, DNA damage levels increased rapidly, peaking between 0.5 and 2 hours, and declined from 4 hours onwards, consistent with active DNA repair (Figure 5F-H and Supplementary Figure 5D). This DNA damage response was accompanied by an acute and progressive increase in H3K27me3 levels, indicating that histone methylation is a rapid chromatin response to DNA damage rather than a delayed epigenetic remodeling event (Figure 5G-H). Inhibition of SAH hydrolysis with NepA exacerbated DNA damage and impaired its resolution, while simultaneously reducing the irradiation induced increase in histone methylation. These findings indicate that methylation activity contributes functionally to DNA damage control in TP53-deficient cells.

To assess how acute DNA damage and inhibition of histone methylation affect ATP availability, ATP levels were measured in whole cell extracts and isolated nuclei (Supplementary Figure 5E-F) following irradiation in the presence or absence of NepA. Cellular and nuclear ATP levels remained stable following DNA damage induction (Figure 5I-J). In contrast, while NepA treatment did not alter whole cell ATP content, it caused a significant decrease in nuclear ATP levels, coinciding with the temporal dynamics of increased DNA damage and reduced H3K27me3 (Figure 5I-J). Together, these data establish a functional link between DNA damage-induced methylation, SAH-derived adenosine production, nuclear ATP synthesis, and efficient DNA repair.

Collectively, our findings support a model in which TP53 loss imposes a chronic demand for nuclear ATP driven by accelerated DNA replication and elevated DNA damage. Under conditions of nucleotide stress or acute DNA damage, this demand triggers increased methylation activity that locally recycles adenosine generates ATP within the nucleus for efficient DNA damage repair. Disruption of this nuclear methylation-adenosine-ATP axis prevents TP53-deficient cells from meeting nuclear energetic demands, ultimately leading to defective genome maintenance.

### TP53 loss creates a tumour specific requirement for adenosine dependent purine metabolism

Our results indicate that loss of TP53 creates a requirement for purine synthesis that is specifically linked to adenosine metabolism and supports tumour cell growth under conditions of elevated cellular stress. Consistent with this model, pharmacological inhibition of de novo purine synthesis in TP53-deficient cells leads to increased DNA damage accumulation and reduced nuclear ATP availability, suggesting that purine metabolism in TP53-deficient cells becomes particularly important because cellular demand for nuclear energy is increased.

We therefore reasoned that inhibition of purine synthesis in TP53-deficient cells would synergise with DNA damage induction, as DNA damage clearance is compromised under conditions of purine limitation. To test this hypothesis, TP53-proficient and TP53-deficient cells were treated with increasing concentrations of 6MP in combination with either paclitaxel, a microtubule stabilizer that primarily perturbs mitotic progression^46^, or etoposide, a topoisomerase II inhibitor that induces DNA double-strand breaks^47^. Whereas co-treatment with paclitaxel resulted in largely additive effects on cell growth (Figure 6A, Supplementary Figure 6A and Supplementary Data 23), the combination of 6MP with etoposide produced a marked synergistic reduction in proliferation, most pronounced in TP53-deficient cells (Figure 6B, Supplementary Figure 6B and Supplementary Data 24). This selective interaction with a DNA-damaging agent indicates that adenosine dependent purine metabolism becomes particularly limiting when cells experience increased genotoxic stress.

**Figure 6.**
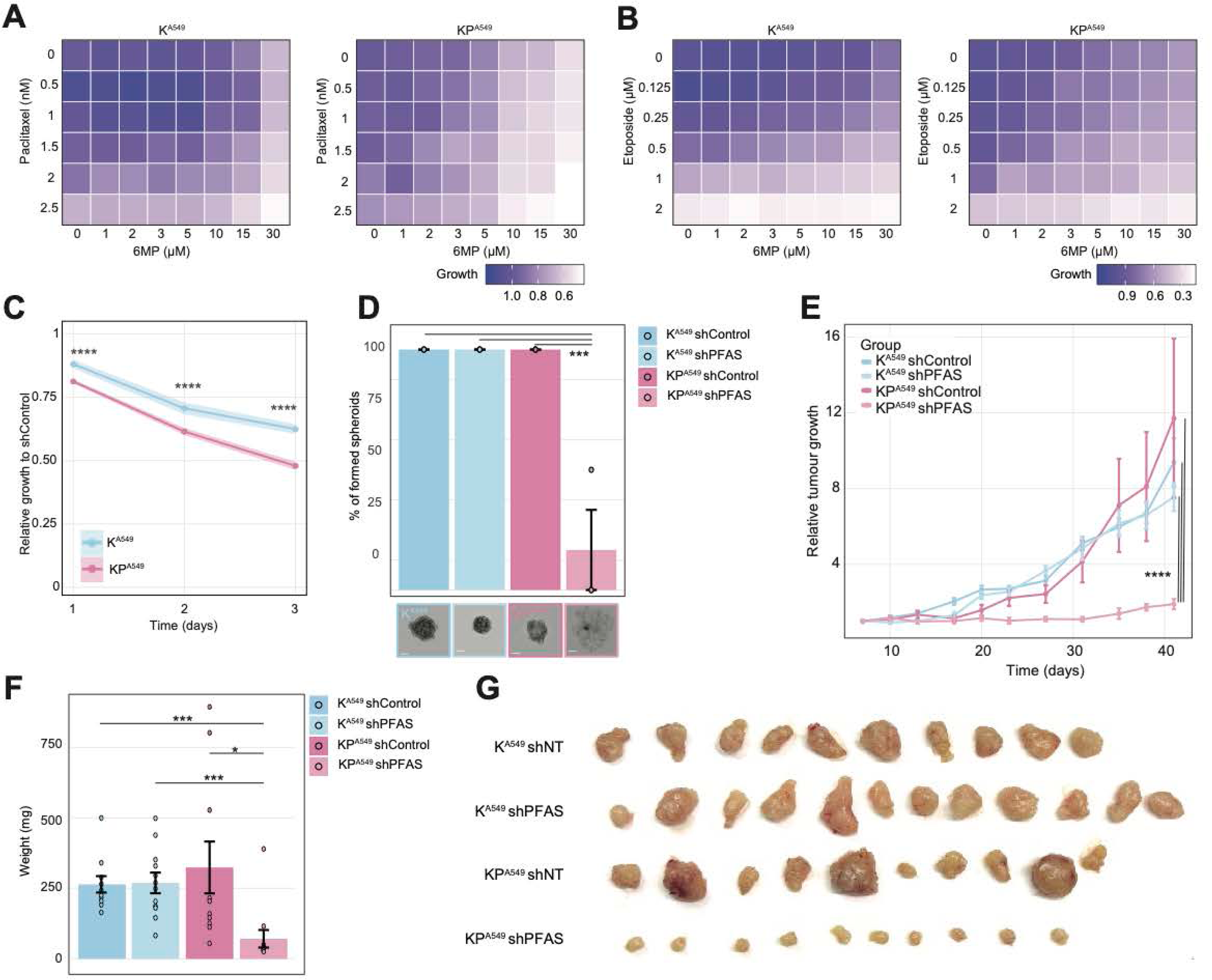
PFAS depletion selectively impairs tumour growth in TP53-deficient cancer cells. Co-treatment matrix for K^A549^ and KP^A549^ treated with 6MP in combination with A) paclitaxel and B) etoposide at the indicated concentrations. Data are normalised to the DMSO-treated condition, following NLS-GFP signal analysis of nuclei count. C) Batch corrected relative cell growth of K^A549^ and KP^A549^ cells transduced with shPFAS, normalised to scrambled (shControl) and analysed by NLS-GFP signal of nuclei count. D) (Top) Quantification of number of K^A549^ and KP^A549^ spheroid formation following transduction with shControl or PFAS shRNAs. (Bottom) representative images. Scale bar = 100µm. N=3, Bonferroni adjusted Wilcoxon pairwise comparison. ***P ≤ 0.0005, ****P ≤ 0.00005. In vivo subcutaneous tumour growth into NSG mice measured over 42 days. E) Tumour volume relative to volume after one week of injection for indicated genotypes. Mean ± SEM. ****P ≤ 0.00005; p-values were calculated with Linear Mixed Effect Model with POST-HOC Tukey HSD correction. F) Tumour weight at ethical endpoint. N = 10-12 tumours/genotype. Bonferroni adjusted Wilcoxon pairwise comparison. ***P ≤ 0.0005.

To genetically validate the requirement for de novo purine synthesis in supporting tumour cell growth under TP53 loss, we next targeted key enzymes within the pathway. Based on our CRISPR-Cas9 screen, we selected GART and PFAS, two central components of *de novo* purine biosynthesis, and depleted their expression using shRNAs in TP53-proficient and TP53-deficient A549 cells (Supplementary Figure 6C). While knockdown of GART impaired proliferation in both genotypes to a similar extent (Supplementary Figure 6D), depletion of PFAS resulted in a pronounced reduction of proliferation specifically in TP53-deficient cells, both in two-dimensional cultures (Figure 6C and Supplementary Data 25) and in three-dimensional spheroid assays (Figure 6D). In contrast, PFAS knockdown had a substantially weaker effect on TP53-proficient cells, indicating a selective requirement for this enzyme upon TP53 loss.

To assess whether this dependency extends to tumour growth *in vivo*, TP53-proficient and TP53-deficient A549 cells expressing either control shRNAs or PFAS-targeting shRNAs were transplanted subcutaneously into immunocompromised mice. Whereas PFAS depletion had little effect on tumour growth in TP53-proficient cells, loss of PFAS strongly impaired the growth of tumours derived from TP53-deficient cells (Figure 6E–G and Supplementary Figure 6E). These findings demonstrate that PFAS-dependent purine synthesis is required to sustain tumour growth in the absence of TP53.

Together, these results establish that TP53 loss creates a tumour specific dependency on adenosine dependent purine metabolism. Rather than reflecting a general requirement for nucleotide supply, this dependency emerges under conditions of elevated cellular stress and is selectively unmasked when purine synthesis is perturbed. These findings place adenosine dependent purine metabolism as a central metabolic adaptation associated with TP53 loss that supports tumour growth.

## Discussion

In this study, we identify a metabolic dependency that emerges upon loss of TP53 and is rooted in the compartmentalized regulation of purine metabolism. By integrating genetic screening, metabolomics, functional perturbations, and *in vivo* models, we show that TP53 deficiency imposes a selective requirement for adenosine dependent purine metabolism that supports cellular fitness under conditions of elevated genotoxic stress. Rather than reflecting a general increase in biosynthetic demand, this dependency reveals how metabolic pathways are reorganised to meet nuclear specific requirements when genome surveillance is compromised.

TP53 is a central regulator of genome integrity and cellular stress responses^3^. Beyond its canonical roles in cell cycle control and apoptosis^48^, TP53 contributes to the coordination of metabolic programs that align energy availability with genome-associated processes^49^. Loss of TP53 disrupts this coordination, leading to elevated baseline DNA damage and persistent activation of DNA repair pathways^10^. Our data extend this framework by showing that TP53 loss is associated with altered purine metabolism that is not explained by differences in proliferation rate. TP53-proficient and TP53-deficient isogenic lung cancer cells proliferate at comparable rates, yet TP53-deficient cells exhibit a selective dependence on de novo purine synthesis and adenosine-related metabolites. This dependency becomes apparent when de novo purine synthesis is inhibited. Pharmacological blockade with 6MP induces a pronounced accumulation of DNA damage and replication stress in TP53-deficient cells, together with a reduction in nuclear ATP levels. Concomitantly, we observe a marked increase in repressive histone methylation marks, including H3K9me3 and H3K27me3, which is substantially stronger in TP53-deficient cells than in TP53-proficient counterparts. Importantly, both the increase in DNA damage and the elevation in histone methylation are reversed by adenosine supplementation, directly linking these phenotypes to adenosine availability rather than to nonspecific toxicity or global nucleotide depletion. These observations indicate that inhibition of purine synthesis imposes acute nuclear metabolic stress in TP53-deficient cells. Reduced adenosine availability can limit nuclear ATP supply, while increased DNA damage further elevates the energetic demands associated with chromatin remodelling and DNA repair. The accompanying increase in histone methylation suggests that chromatin responds actively to changes in nuclear metabolic state. Indeed, a central implication of our findings is that chromatin itself functions as a dynamic metabolic interface. Histone methylation is not only a regulatory modification that shapes chromatin states, but also a metabolically demanding process that consumes SAM. In TP53-deficient cells, increased histone methylation upon purine synthesis inhibition coincides with heightened dependence on adenosine availability and is reversed by adenosine supplementation, directly linking chromatin modification to nuclear metabolic state. Histone methylation also increases rapidly following acute DNA damage induced by low-dose γ-irradiation that is fully recoverable, indicating that this response is engaged early and does not reflect delayed epigenetic remodelling. Together, these observations suggest that histone methylation functions as an active, metabolism-associated chromatin response to nuclear stress rather than as a secondary adaptation to DNA damage.

We propose that, under conditions of elevated nuclear stress, increased histone methylation promotes the conversion of SAM to SAH, which is enzymatically cleaved by AHCY to homocysteine and adenosine. Through this AHCY-dependent reaction, chromatin-associated methylation becomes functionally coupled to SAM–adenosine recycling, contributing to the maintenance of nuclear adenosine and ATP availability required to cope with DNA damage. Support for this model comes from the increased sensitivity of TP53-deficient cells to AHCY inhibition, indicating that efficient enzymatic processing of SAH is functionally important in this context. Importantly, this interpretation reframes the observed increase in histone methylation not as a secondary epigenetic consequence of metabolic stress, but as part of an adaptive metabolic response that links chromatin modification to nuclear energy homeostasis. In this view, chromatin acts as a metabolic buffer that can transiently incorporate methyl groups through histone methylation while remaining functionally coupled to pathways that replenish adenosine and sustain nuclear ATP levels.

The interaction between purine synthesis inhibition and DNA-damaging agents further supports the idea that this dependency is revealed under conditions of heightened nuclear stress. Synergistic effects observed upon co-treatment with etoposide, but not with agents that primarily perturb mitosis, indicate that adenosine-dependent purine metabolism becomes particularly limiting when DNA repair demands are increased. These findings reinforce the notion that the metabolic vulnerability uncovered here is functionally linked to nuclear stress rather than to cell division alone.

Finally, genetic depletion of PFAS selectively impairs the growth of TP53-deficient tumours in vivo, demonstrating that this metabolic requirement extends beyond cultured cells. While in vivo tumour growth integrates multiple layers of stress, these results support the idea that adenosine-dependent purine metabolism represents a critical adaptive pathway that sustains cellular fitness when TP53-mediated control is absent.

Together, our findings reposition purine metabolism as a spatially and functionally organized process that supports nuclear adaptation to genotoxic stress. Rather than acting solely as a source of nucleotides for proliferation, adenosine-dependent purine metabolism emerges as a compartmentalized metabolic program that links chromatin regulation to nuclear energy balance. More broadly, this work highlights how loss of genome surveillance can unmask latent dependencies on nuclear metabolic regulation, providing a framework for understanding how metabolism, chromatin state, and genome stability are integrated at the subcellular level.

## Methods

### Cell Culture

All cell lines were maintained under standard tissue culture conditions at 37°C in a humidified incubator with 5% CO₂ and were passed when they reached 80% confluency. A549 human lung adenocarcinoma cells were cultured in Dulbecco’s Modified Eagle Medium (DMEM, Labclinics, #L0102-500) supplemented with 10% foetal bovine serum (FBS, ThermoFisher, #A5256701) and 1% Penicillin-Streptomycin (PenStrep, Life Technologies, #15140122). HBEC3-KT cells, derived from immortalised human bronchial epithelial cells, were cultured in Roswell Park Memorial Institute (RPMI) medium (Thermo Scientific, #11879020) also supplemented with 10% FBS and 1% PenStrep. KPC270^PDAC^ and KPC172^PDAC^ cells were grown in DMEM containing 10% FBS and 1% PenStrep. M1196 KPshp53 cells (KC^PDAC^), which harbour a doxycycline-inducible shRNA targeting Trp53, were grown on collagen-coated plates (PurCol, Advanced Biomatrix; final concentration 0.1 mg/mL, Corning® Collagen, #CLS354236). They were maintained in DMEM with 10% FBS, 1% PenStrep, and 1 μg/mL doxycycline to maintain Trp53 knockdown (KPC^PDAC^). Doxycycline was removed from the medium to restore Trp53 expression, and complete withdrawal was ensured by washing cells three times with pre-warmed phosphate-buffered saline (PBS) followed by replacement with DMEM containing 10% tetracycline-free FBS and PenStrep. Trp53 re-expression was confirmed after 72 hours post-doxycycline withdrawal. Cells engineered with doxycycline-inducible TP53 expression constructs were handled similarly, with doxycycline (1 µg/mL) refreshed every 2–3 days to maintain gene induction. For initial expansion after thawing, cells were cultured with doxycycline and switched to tetracycline-free conditions for induction once optimal confluency was achieved. Cells were routinely frozen in FBS with 10% dimethyl sulfoxide (DMSO) and stored at −80°C.

### Lentiviral Production

HEK 293T cells were used for lentiviral packaging of plKO.1 and maintained in DMEM supplemented with 10% FBS and 1% PenStrep. For viral production, 15 × 10⁶ 293T cells were seeded into 15-cm tissue culture dishes containing 20 mL of complete DMEM. Once the cells reached approximately 80% confluency on the following day, transfection was carried out using polyethylenimine (PEI) as the transfection reagent. Between 6 and 8 hours after transfection, the medium was replaced with 20 mL of fresh DMEM. Forty-eight hours after transfection, viral supernatants were collected, filtered through a 0.45 µm polyethersulfone (PES) membrane to remove cellular debris, and aliquoted into 2 mL volumes before being stored at −80°C for future use.

### Human Metabolic CRISPR Screen

The pooled CRISPR screen targeting 2,981 human metabolic genes was performed using a lentiviral sgRNA library obtained from Addgene (Pooled Library #110066), which includes 29,790 guide RNAs. Library amplification was carried out using the QIAGEN Plasmid Plus Mega Kit (Cat. No. 12981). A549 TP53 proficient and deficient cells were transduced at a multiplicity of infection (MOI) of 0.3 to ensure single-copy integration and a 1000× coverage of the sgRNA library. After 24 hours, puromycin was added at a concentration of 2.5 µg/µL to select for transduced cells. Selection continued for 48 hours, after which cell pellets were collected for baseline (Day 0) analysis.

Post-selection, cells were cultured for an additional 14 days while maintaining sufficient coverage of the library. On Day 14, final cell pellets were harvested for downstream analysis. Genomic DNA was extracted from both Day 0 and Day 14 samples using the QIAGEN QIAamp DNA Blood Mini Kit (Cat. No. 250). A two-step PCR protocol was employed to amplify the sgRNA regions and prepare them for next-generation sequencing (NGS).

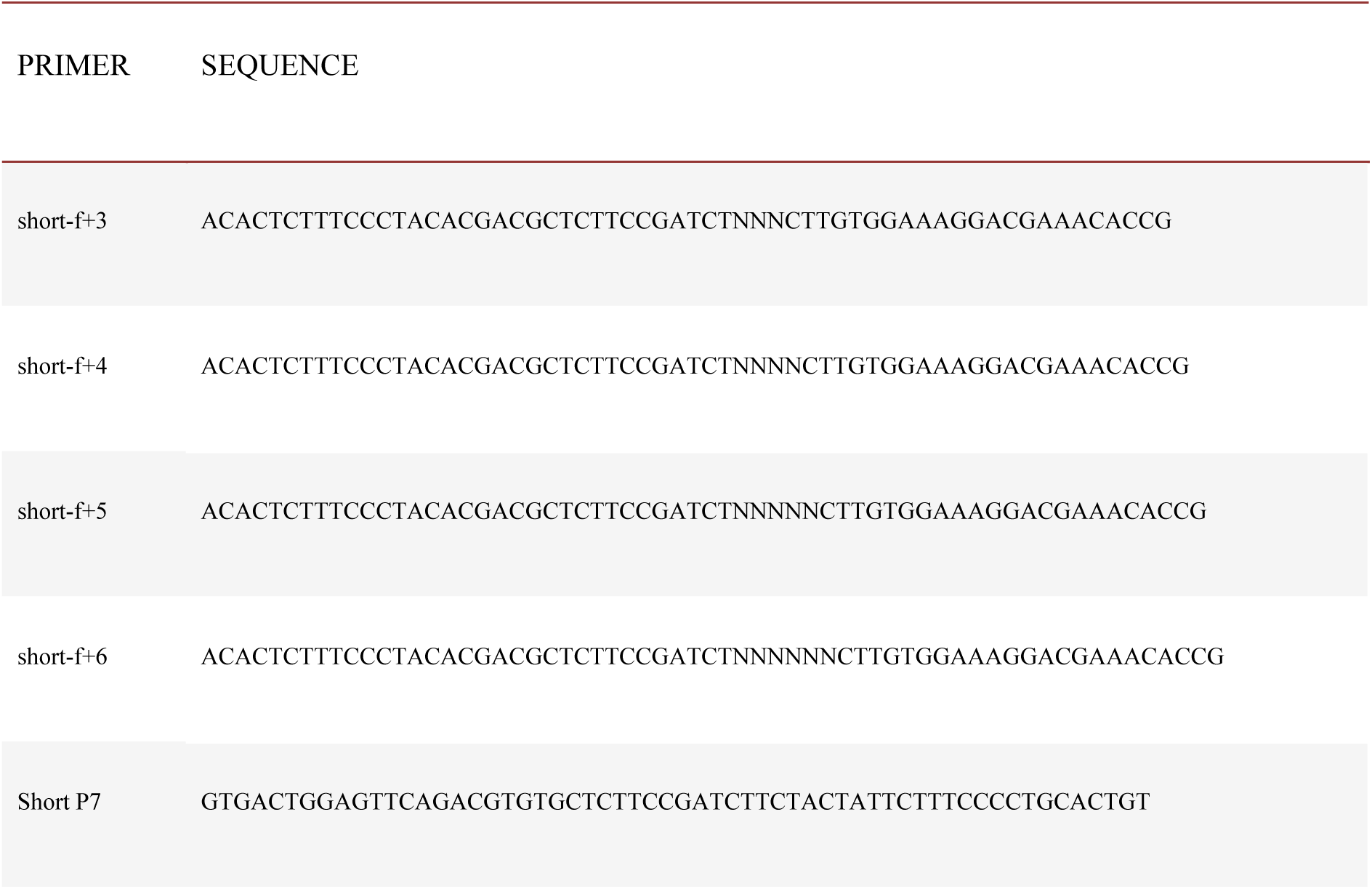

Libraries were sequenced at the CRG Genomics Facility using Illumina platforms. MAGeCK (Model-based Analysis of Genome-wide CRISPR/Cas9 Knockout) software was used to identify genes whose loss was differentially enriched or depleted in TP53 knockout versus control cells.

### Drug Sensitivity and Proliferation Assays

Live-cell proliferation and drug response were assessed in A549 and HBEC3-KT cells expressing Cas9 and either non-targeting (sgNT) or TP53-targeting (sgTP53) sgRNAs. Cells were infected with the pTRIP-SFFV-EGFP-NLS lentiviral construct (Addgene #86677) and sorted for GFP-positive cells. A total of 2,500 A549 or 3,000 HBEC3-KT cells were seeded per well in 96-well Phenoplate black-bottom plates (PerkinElmer). The following day, cells were treated with increasing concentrations of various drugs including Methotrexate (MedChem Express, HY-14519), Neplanocin A (Cayman Chemical, 10584) and 6MP (MedChem Express, HY-13677). For combination therapy experiments, A549 cells were treated with increasing doses of 6MP in combination with increasing doses of Etoposide (MedChem Express, #HY-13629) or Paclitaxel (MedChem Express, #HY-B0015). Drug exposure continued for 72 hours, and cell proliferation was monitored using the Operetta high-content imaging system (PerkinElmer) and analysed using the Harmony software (version 4.9).

### Drug Treatment in Doxycycline-Inducible Cells

For doxycycline dependent TP53 induction experiments, KC^PDAC^ pancreatic cells were seeded and cultured with or without doxycycline. After 24 hours, cells were reseeded into 96-well Operetta-compatible plates at 4,000 cells per well and treated with increasing doses of 6MP. Hoechst dye (Thermo Fisher, #H3570) was added at time points 0 and 72 hours to stain cell nuclei, and imaging was performed using the Operetta high-content imaging system (PerkinElmer) and analysed using the Harmony software (version 4.9). Protein samples were collected in parallel for Western blot analysis.

### Rescue Proliferation Assays with Metabolites

To assess rescue of drug-induced cytotoxicity, A549 and HBEC3-KT cells expressing sgNT or sgTP53 were treated with a fixed dose of 6MP alongside increasing concentrations of various metabolites. Tested compounds included S-adenosylmethionine (Sigma-Aldrich, #A4377), S-adenosylhomocysteine (Sigma-Aldrich, #A9384), NAD⁺ (Thermo Fisher, #J62337.03), FAD (Sigma-Aldrich, #F8384), Adenosine 5’-triphosphate (Thermo Fisher, #R0441), Guanosine 5’-triphosphate (Thermo Fisher, #R0461), Adenosine (Sigma-Aldrich, #A4036), Guanosine (Sigma-Aldrich, #G6264), Hypoxanthine (Sigma-Aldrich, #H9636), Cytidine 5’-triphosphate (Thermo Fisher, #R0451), and Uridine 5’-triphosphate (Thermo Fisher, #R0471). Proliferation was measured after 72 hours using the Operetta high-content imaging system (PerkinElmer) and analysed using the Harmony software (version 4.9).

### Spheroids experiments

1,000 A549 cells were seeded in Ultra-low attachment 96-well plates (CellCarrier Spheroid ULA 96-well Microplates, PerkinElmer, #6055330) and centrifuged at 1,000 rpm for 10 minutes. The corresponding treatment was added with the seeding, being that increasing doses of 6MP with or without the supplementation of adenosine. Images of the spheroids were taken a week after seeding with the Operetta high-content imaging system (PerkinElmer), and the area of the spheroids was quantified using ImageJ.

### DNA Fiber Assay

Replication fork dynamics were analysed in TP53 deficient and proficient HBEC3-KT cells. Cells were seeded at a density of 1.25 × 10⁵ cells per 6-cm dish and allowed to adhere for 96 hours. Cells were treated with 5 µM 6MP or DMSO for 72 hours prior to labelling. DNA replication was tracked using sequential pulse labelling with thymidine analogues. Cells were first incubated with 25 µM chlorodeoxyuridine (CldU, Sigma-Aldrich, #C6891) for 20 minutes at 37°C, followed by a wash and a second 20-minute incubation with 250 µM iododeoxyuridine (IdU, Sigma-Aldrich, #I7125). After labelling, cells were washed twice with ice-cold PBS, trypsinized, and resuspended in cold PBS at a concentration of 1 × 10⁶ cells/mL. For DNA fibre spreading, 2 µL of the cell suspension was placed onto a clean microscope slide, allowed to dry partially for 5–7 minutes, and mixed gently with 7 µL of DNA spreading buffer (200 mM Tris-HCl pH 7.4, 50 mM EDTA, 0.5% SDS). The slide was tilted to allow fibre spreading by gravity, air-dried, and fixed in methanol:acetic acid (3:1) for 10 minutes before storage at 4°C. For immunodetection, DNA was denatured by incubating slides in 2.5 M HCl for 1 hour at room temperature. Slides were then blocked with PBS + 1% BSA + 0.1% Tween-20 for 1 hour. Then, slides were incubated with rat anti-BrdU (Abcam, #ab6326) and mouse anti-BrdU (BD Biosciences, #347580), each diluted 1:500 in blocking buffer, for 1.5 hours at 37°C. After PBS washes, slides were incubated with Alexa Fluor 488-conjugated anti-rat (Invitrogen, #A-11006) and Alexa Fluor 555-conjugated anti-mouse (Invitrogen, #A-21424) secondary antibodies (1:500 dilution). Samples were mounted using Vectashield and imaged with a 63x objective via confocal microscopy.

### Metabolomics

For metabolite profiling, HBEC3-KT cells were seeded in 6-well plates 96 hours before extraction to reach a confluency of 750,000 to 1,000,000 cells per well. Cells were treated for 72 hours with either 5 µM 6MP or DMSO in a medium containing dialyzed FBS (Thermo Fiser, #A3382001). Two hours before extraction, fresh medium was added to promote a steady-state metabolic condition. All extractions were performed on ice. Cells were washed three times with ice-cold blood bank saline, and residual liquid was carefully aspirated. Two hours before extraction, media on the cells was exchanged with fresh media. To extract metabolites, cells were placed on iced, washed three times with ice-cold blood bank saline, and 500 µL of ice-cold 80% methanol (HPLC grade) with 500 nM 13C/15N-labelled amino acid standards was added to each well. Cells were thoroughly scraped, collected, and transferred into chilled microcentrifuge tubes. Extracts were immediately frozen at −80°C and later shipped on dry ice for metabolomic analysis.

Samples were vortexed for 10 min at 4°C, centrifuged at maximum speed for 10 min at 4°C after which the supernatant was collected. Samples were dried under a gentle stream of nitrogen and then reconstituted in 25 µL of a 50:50 (v/v) mixture of HPLC-grade acetonitrile and water. Metabolite analysis was performed on a Dionex UltiMate 3000 ultra-high performance liquid chromatography system (Thermo Fisher Scientific) coupled to a Q Exactive Orbitrap mass spectrometer equipped with an Ion Max source and HESI II probe. The instrument was externally calibrated every seven days using the manufacturer’s standard calibration mix. For chromatographic separation, 2 µL of each sample was injected onto a SeQuant ZIC-pHILIC polymeric column (2.1 × 150 mm, 5 µm particle size; Millipore Sigma). The column compartment and autosampler tray were maintained at 25 °C and 4 °C, respectively, with a flow rate of 150 µL/min. Mobile phase A was 20 mM ammonium carbonate containing 0.1% ammonium hydroxide, and mobile phase B consisted of 100% acetonitrile. The mobile gradient program was 0–20 min, linear transition from 80% to 20% B; 20–20.5 min, rapid return to 80% B; 20.5–28 min, hold at 80% B. Ionization parameters were: sheath gas 40, auxiliary gas 15, sweep gas 1, spray voltage 3.1 kV, capillary temperature 275 °C, probe heater 350 °C, and S-lens RF level 40. Data were collected in full-scan mode with polarity switching across an m/z range of 70–1000. To improve sensitivity for nucleotides, an additional negative-mode full scan from m/z 220–700 was acquired. Mass spectrometer resolution was set to 70,000, with an AGC target of 1 × 10^6 and a maximum injection time of 20 ms.

Metabolite levels were quantified in XCalibur, applying a 5-ppm mass tolerance. Raw peak areas of metabolites were normalized by cell count for each sample and to the abundance of internal 13C/15N-labelled amino acid standards.

After determining the normalized peak area for metabolite in each sample, one factor statistical analysis was performed on the data using Metaboanalyst 6.0. Metabolites with more than 50% missing values were excluded from further analysis, while for the remaining metabolites, missing entries were imputed as one-fifth of the lowest detected positive value. Auto scaling was used.

### Flow Cytometric Analysis of Cell Cycle

TP53 deficient and proficient A549 cells were seeded in 12-well plates at a density of 30,000 cells per well and treated for 72 hours with 15 µM 6MP or DMSO as a control. 10 µM EdU was added for 2 hours prior to harvesting to label actively proliferating cells. Cells were then collected, centrifuged at 1,500 rpm for 5 minutes, and washed in PBS with 1% BSA.

### shRNA-Mediated Knockdown

Short hairpin RNAs (shRNAs) targeting PFAS and GART were cloned into the pLKO.1-TRC vector by removing the stuffer sequence with EcoRI and AgeI digestion and ligating annealed oligonucleotides. Oligos were annealed by heating to 95°C followed by slow cooling. The digested vector was gel purified and ligated with shRNA duplexes using NEB T4 DNA ligase. Ligations were transformed into chemically competent E. coli and plated on LB agar with 100 µg/mL ampicillin. Colonies were picked and sequenced to confirm correct insertion. Lentivirus was generated using HEK293T cells as described previously.

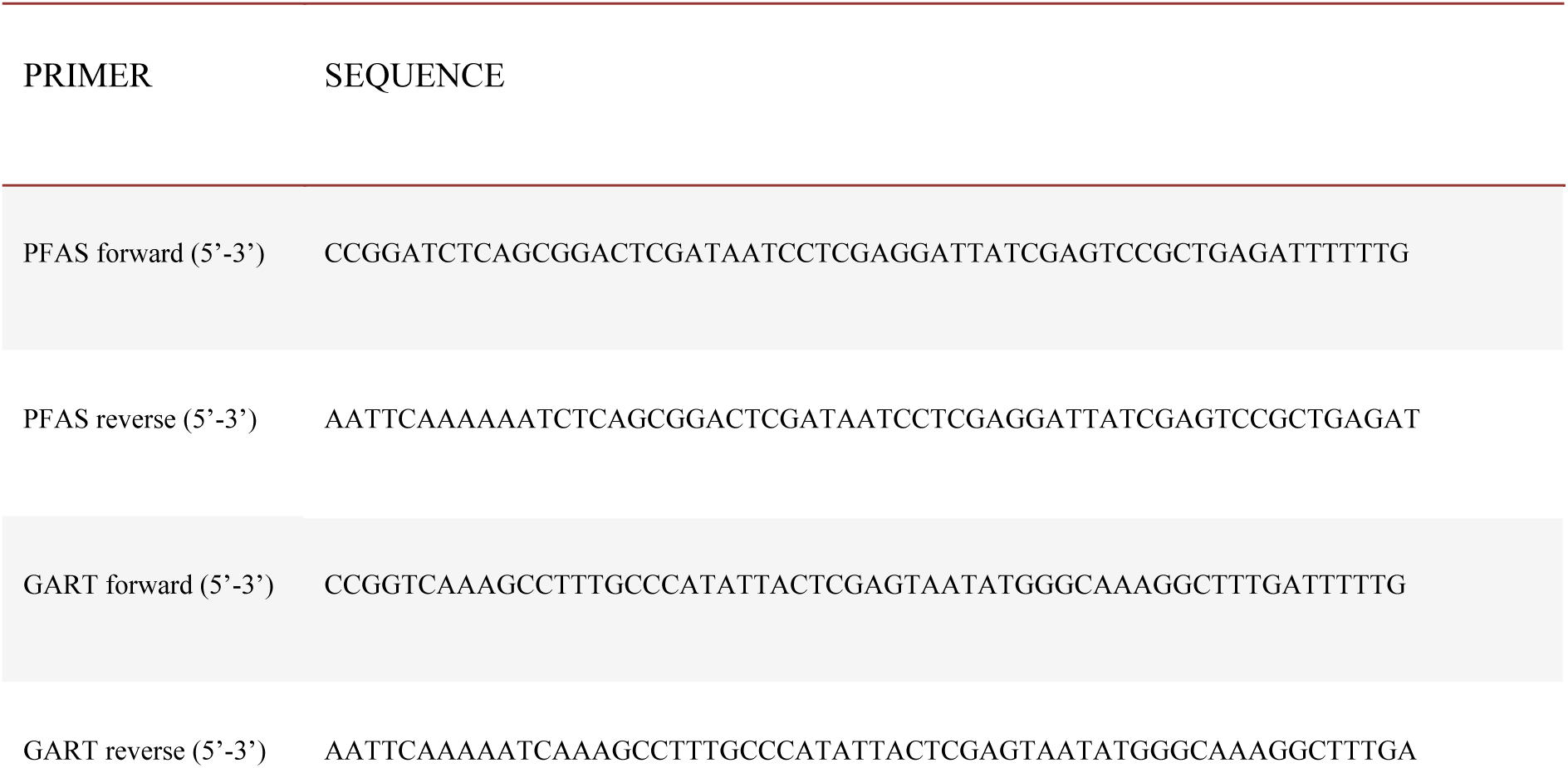

### Quantitative Real-Time PCR (qRT-PCR)

Total RNA was extracted from cultured cells using TRIzol™ Reagent (Thermo Fisher Scientific, #15596018), and 1 µg of total RNA was reverse transcribed using SuperScript™ IV Reverse Transcriptase and oligo(dT) primers (Thermo Fisher Scientific, #18090050). Quantitative PCR was performed with the LightCycler® 480 SYBR Green I Master Mix (Roche, #04887352001) on a QuantStudio™ 12K Flex Real-Time PCR System. The relative expression of target genes was calculated using the ΔΔCt method and normalized to GAPDH. All samples were analysed in biological triplicates and technical duplicates.

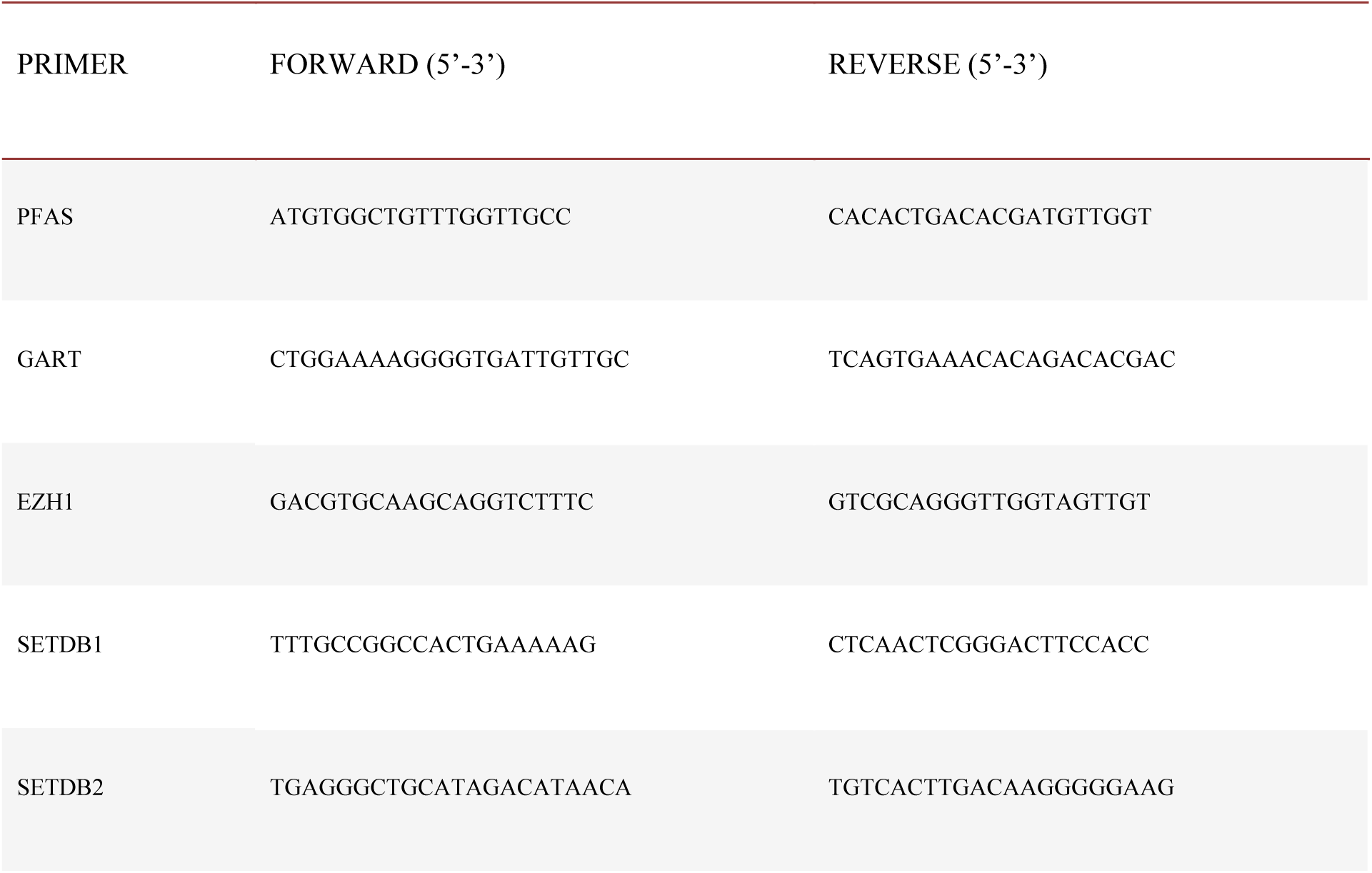

### Nuclei extraction

1 million TP53 deficient A549 cells were seeded into a 6-well plate and after 24 hours, the cells were treated with 10 µM Neplanocin A or DMSO and γ-irradiated or not with 2 Gy. Nuclei were isolated using a rapid hypotonic lysis protocol and all steps were performed on ice. Cells were harvested by scraping in a hypotonic buffer (10 mM Tris-HCl pH 7.4, 10 mM KCl, 2 mM MgCl₂, 0.25 M sucrose) and centrifuged at 600 g for 3 min at 4°C. Cell pellets were resuspended in the hypotonic buffer containing 0.1% NP-40 and incubated on ice for 4 min. Nuclei were collected by centrifugation at 600 g for 4 min at 4°C and washed once in isotonic wash buffer (10 mM Tris-HCl pH 7.4, 150 mM KCl, 2 mM MgCl₂, 0.25 M sucrose). Nuclei were counted using Trypan blue staining and an automatic cell counter Countess 3.

### Immunofluorescence

For immunostaining, cells were seeded in 96-well plates (CellCarrier-96 Ultra Microplates, PerkinElmer, #6055302), allowed to adhere overnight and then treatment was added. The treatments include 6MP or DMSO with or without supplementation of 100 µM adenosine for 72 hours, or 10 µM of Neplanocin A or DMSO for a time course of maximum 6 hours with and without γ-irradiation of 2 Gy. After the treatment, cells were fixed in 4% paraformaldehyde for 10 minutes at room temperature. Then, cells were washed with PBS, permeabilized, and blocked with PBS containing 5% BSA and 0.3% Triton X-100 for 1 hour. Primary antibodies including γH2AX (Cell Signaling Technology, #9718S, 1:2000; and Sigma-Aldrich, #05-636-I, 1:500), 53BP1 (Novus, #NB100-304SS, 1:5000 and BioLegend, #933002, 1:500), H3K9me3 (Diagenode, C15410193, 1:500) and H3K27me3 (Diagenode, #C15410195, 1:500) were diluted in the same buffer and incubated overnight at 4°C.Following primary incubation, wells were washed and incubated for 10 minutes at room temperature with Alexa Fluor-conjugated secondary antibodies Alexa 647 anti-rabbit (Invitrogen, #A21244), Alexa 647 anti-rat (Abcam, #ab150167, 1:500), Alexa 555 anti-mouse (Invitrogen, #A21424, 1:500) and Alexa 405 anti-rabbit (Abcam, #ab175649, 1:500). Nuclei were counterstained with DAPI (1 µg/mL) for 10 minutes. Plates were imaged using the Operetta high-content imaging system (PerkinElmer) and analysed using the Harmony software (version 4.9).

### ATP quantification

ATP was quantified using the ATP Determination Kit (Invitrogen™, #A22066). Nuclear pellets and matched whole-cell pellets were lysed with a lysis buffer (10 mM Tris-HCl pH 7.4, 100 mM NaCl, 1mM EDTA, 0.01% Triton X-100). ATP standards were prepared fresh in the same lysis buffer and used to generate a standard curve on each plate. Reactions were assembled in white 96-well plates (Thermo Fisher Scientific #136101) by adding 90 µL of the ATP reaction mix and 10 µL of the nuclear or whole cell lysates. After a 15 minute incubation, the luminescence was recorded on a TECAN Infinite M200 Plate Reader, and ATP concentrations were interpolated from the standard curve with background subtraction.

### Western Blotting

Cells were collected and lysed in ice-cold RIPA buffer (Thermo Fisher Scientific) supplemented with protease inhibitors (Roche, cOmplete™) to obtain whole-cell extracts. Nuclei were extracted following the protocol described in the *Nuclei extraction* section to obtain nuclei extracts. Protein concentration was quantified using the Pierce BCA Protein Kit (Thermo Fisher, #23225). Equal amounts of proteins were separated by sodium dodecyl sulfate-polyacrylamide gel electrophoresis (SDS–PAGE) method and transferred onto nitrocellulose membranes (Amersham, #10600002). Membranes were blocked in 5% non-fat milk in PBS-Tween 0.05% (Tween-20, Sigma-Aldrich, #P9416) for 1 hour at room temperature and incubated overnight at 4°C with primary antibodies against TP53 (Santa Cruz, sc-126), p21 (Abcam, ab188224), β-actin (Santa Cruz, sc-47778), vinculin (E1E9V) (Cell Signaling Technology #13901; 1:1000), Histone H3 (Cell Signalling Technology #14269; 1:10000) and Ferredoxin 1 (FDX1, Thermo Fisher Scientific, #PA5-59653; 1:1000). After washing, membranes were incubated with HRP-conjugated secondary antibody and visualized using ECL Prime Western Blotting Detection Reagent (Cytiva, #RPN2232) on a ChemiDoc MP system (Bio-Rad); or with fluorescence-conjugated secondary antibodies Alexa Fluor 800 goat anti-rabbit IgG (Thermo Fisher, #A32735; 1:10000) and Alexa Fluor 680 goat anti-mouse IgG (Thermo Fisher, #A21058; 1:10000) and visualized with Odyssey CLx Imaging System (LI-COR Biosciences).

### ATP sensor

A total of 5,000 A549 cells were seeded in 96-well Operetta plates and transfected for 30 h with Lipofectamine 2000 (ThermoFisher Scientific, #11668027) according to the manufacturer’s protocol. Cells received either 0.05 ng of pCMV-Cyto luc or Nuclear Luc ICTTO plasmid. Following transfection, cells were treated with 15 µM 6MP or DMSO for 24 hours. Plates were imaged using the Operetta High-Content Imaging System (PerkinElmer) and analysed with Image J.

### Chromatin abundance of enzymes

Chromatin enrichment residuals were obtained from our published chromatome dataset^45^, where protein intensities are corrected by robust linear regression against sample-specific nuclear/chromatin enrichment to derive residual values reflecting relative chromatin association independent of fractionation efficiency. Proteins were restricted to a curated list of one-carbon and methylation-related enzymes, and filtering criteria were relaxed relative to the original study by requiring quantification in at least five samples, after which group-level means were recalculated. Sample metadata, including manually annotated TP53 mutational status, were harmonized into WT and MUT groups (samples annotated as “Unknown” were excluded from group contrasts). A gene-by-sample matrix of chromatin enrichment residuals was z-scored per gene and visualized as a heatmap with samples split by TP53 status and hierarchically clustered within groups using a diverging color scale centered at zero. Group-level differences were computed by averaging chromatin enrichment residuals within WT and MUT samples for each enzyme and calculating the contrast as MUT − WT. The same MUT − WT contrast values were used to color enzyme nodes in pathway schematics, ensuring consistent representation of direction and magnitude across quantitative and pathway-level visualizations.

### In vivo experiments

All animal experiments were conducted in compliance with institutional and national ethical regulations and were approved by the Ethics Committee for Animal Experiments (CEEA-PRBB, Barcelona, Spain). Animals were monitored regularly and euthanized upon reaching predefined humane endpoints, including maximum allowable tumour size or signs of distress. Mice were housed in groups of 5 per cage and irradiated chow and water were provided ad libitum. Subcutaneous tumour models were performed by injection of 1 million A549 cells suspended in a 1:1 mixture of PBS and Matrigel in one flank of 7–10-week-old NSG mice (Envigo). Subcutaneous tumour growth was followed by calliper measurements and the following formula applied to measure tumour volume: volume = 1/2(length × width²). Tumour measurements were performed in a blinded manner.

### Bioinformatic Analysis of Essentialities

Gene expression and mutation data were retrieved from the Cancer Cell Line Encyclopaedia (CCLE). Cell lines were grouped based on TP53 and KRAS mutational status. TP53 pathway activity scores were calculated using the PROGENy algorithm, which infers pathway activation from transcriptomic data. Essentiality scores were obtained from the DepMap Achilles CRISPR dataset and analysed to identify vulnerabilities correlated with TP53 mutation status. All analyses were conducted using R and custom scripts, with visualizations generated using pheatmap and ggplot2.

### Bioinformatic Analysis of TCGA

TCGA RNA-seq and clinical annotations were downloaded from the UCSC Xena browser. Patient barcodes were harmonized (hyphen/period normalization, whitespace trimming) and expression/clinical tables were merged by patient ID. Samples were classified as KRAS/TP53 mutant versus wild-type, considering function-disrupting variants. The tumour stage was standardized by collapsing AJCC pathologic tumour stage (and clinical stage labels) into Stage I–IV, and cohorts were subset by stage (I–III) for downstream analyses. Group sizes by mutational status were tabulated, and stage-specific datasets were exported. All steps were implemented in R using data.table, dplyr, readr, and ggplot2.

## Data availability

All data relevant for the conclusions of this work have been made available through the main text or the supplementary information. All supplementary data have been deposited at: https://zenodo.org/records/16993114?token=eyJhbGciOiJIUzUxMiIsImlhdCI6MTc1NjQ2MjcxNCwiZXhwIjoxNzYxOTU1MTk5fQ.eyJpZCI6IjA1ZjlhNzRkLTIwMTQtNGIxMS1hNjA2LWQ2Y2MzZGE3MDY5MyIsImRhdGEiOnt9LCJyYW5kb20iOiJkOWVkNmRkZDhiZTliYjdmN2EyOTlkMTE0NjFiMmY5NSJ9.A-sfjVh54YvGlLCkKV6EeD_t_zSZLUB76PHu4eQrF9Wo_lKGQcaGWQYduayICHE0EMHc9DwgBS06_clylvfzkQ.

## Code availability

All codes used to generate figures relevant to the metabolomics analysis, as well as other figures derived from source data in this manuscript, can be found at https://github.com/SdelciLab/Urgel-Solas-et-al.-.git.

## Supplementary Figure Legends

**Supplementary Figure 1.**
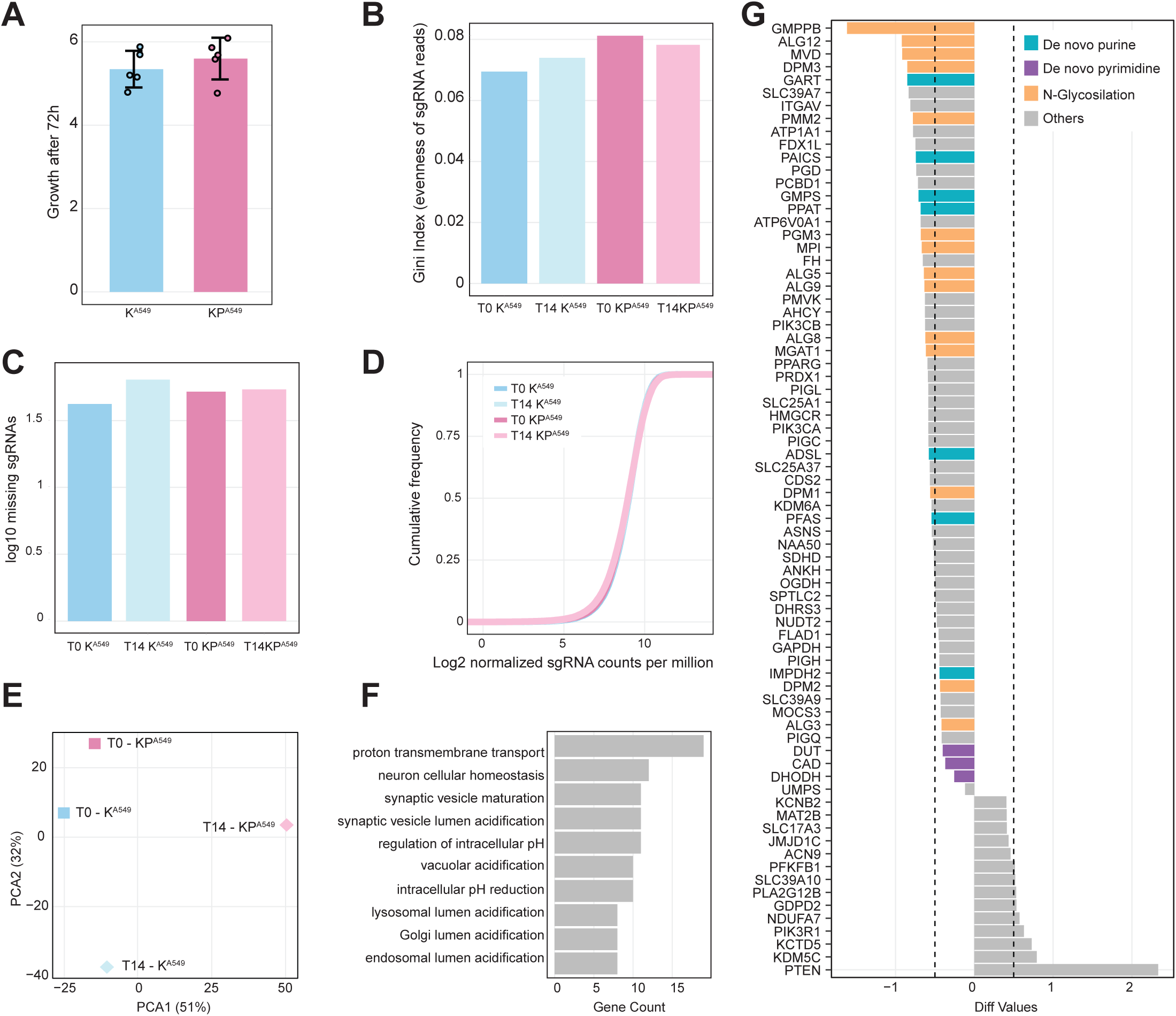
Metabolism-focused CRISPR-Cas9 screen in vitro reveals metabolic dependencies of TP53-deficient cells. A) Relative basal growth of K^A549^ and KP^A549^ after 72h analysed by HT-IF for nuclei counting. Data represents N=5 intendent experiments. Mean ± SD. Two-tailed unpaired t-test. B) Gini index values indicating the evenness of sgRNA read distributions across all conditions: Day 0 and Day 14 for both K^A549^ and KP^A549^ cell lines. C) Log10 number of missing sgRNAs for each condition. D) Cumulative frequency plots of log2-normalized sgRNA counts per million for all four conditions. E) PCA of sgRNA abundance profiles. PC1 (51%) and PC2 (32%) separate samples by cell line and timepoint. F) GO analysis of sgRNAs commonly depleted in K^A549^ and KP^A549^ cell lines. G) Differential dependency analysis between K^A549^ and KP^A549^ cell lines in which genes are ranked by differential log2-fold change (Diff Values) between day 14 and day 0 in both conditions. Genes are annotated by pathway involvement: de novo purine biosynthesis (blue), de novo pyrimidine biosynthesis (purple), N-glycosylation (orange), and others (grey). Negative values represent higher dependency in KP^A549^ cell lines.

**Supplementary Figure 2.**
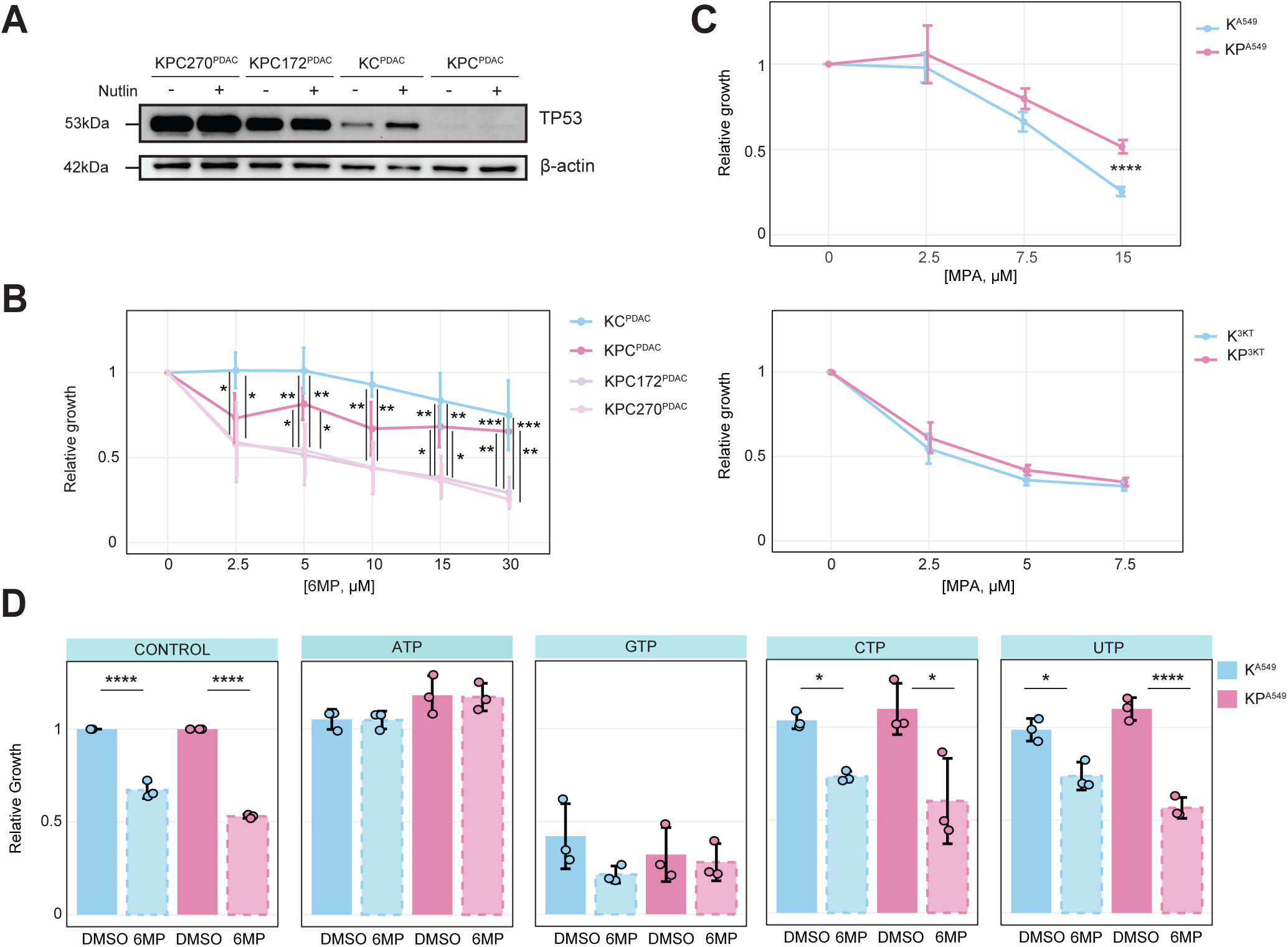
TP53-deficient and TP53-proficient cancer cells respond differently to purine synthesis inhibitors. A) Western blot analysis assessing protein expression of TP53 in the PDAC cell lines. ß-actine was used as a loading control. B) Relative growth of PDAC cell lines after 72 hours treatment with the indicated concentrations of 6MP, normalised to DMSO, and analysed by NLS-GFP signal HT-IF for nuclei counting. Data represents N=4 independent experiments. Mean ± SD. Pairwise t-test. ****P ≤ 0.00005; ***P ≤ 0.0005; **P ≤ 0.005; *P ≤ 0.05. C) Relative growth of K^A549^ and KP^A549^ cells after 72 hours treatment with the indicated concentrations of MPA, normalised to DMSO, and analysed by NLS-GFP signal HT-IF for nuclei counting. D) Relative growth of K^A549^ and KP^A549^ cells treated with 15 µM 6MP and supplemented with 100 µM ATP, GTP, CTP or UTP for 72 hours, normalised to DMSO, and analysed by HT-IF for nuclei counting. Data represents N=3 independent experiments. Mean ± SD. Pairwise t-test with Bonferroni adjustment. P-values ****P ≤ 0.00005; ***P ≤ 0.0005; **P ≤ 0.005; *P ≤ 0.05.

**Supplementary Figure 3.**
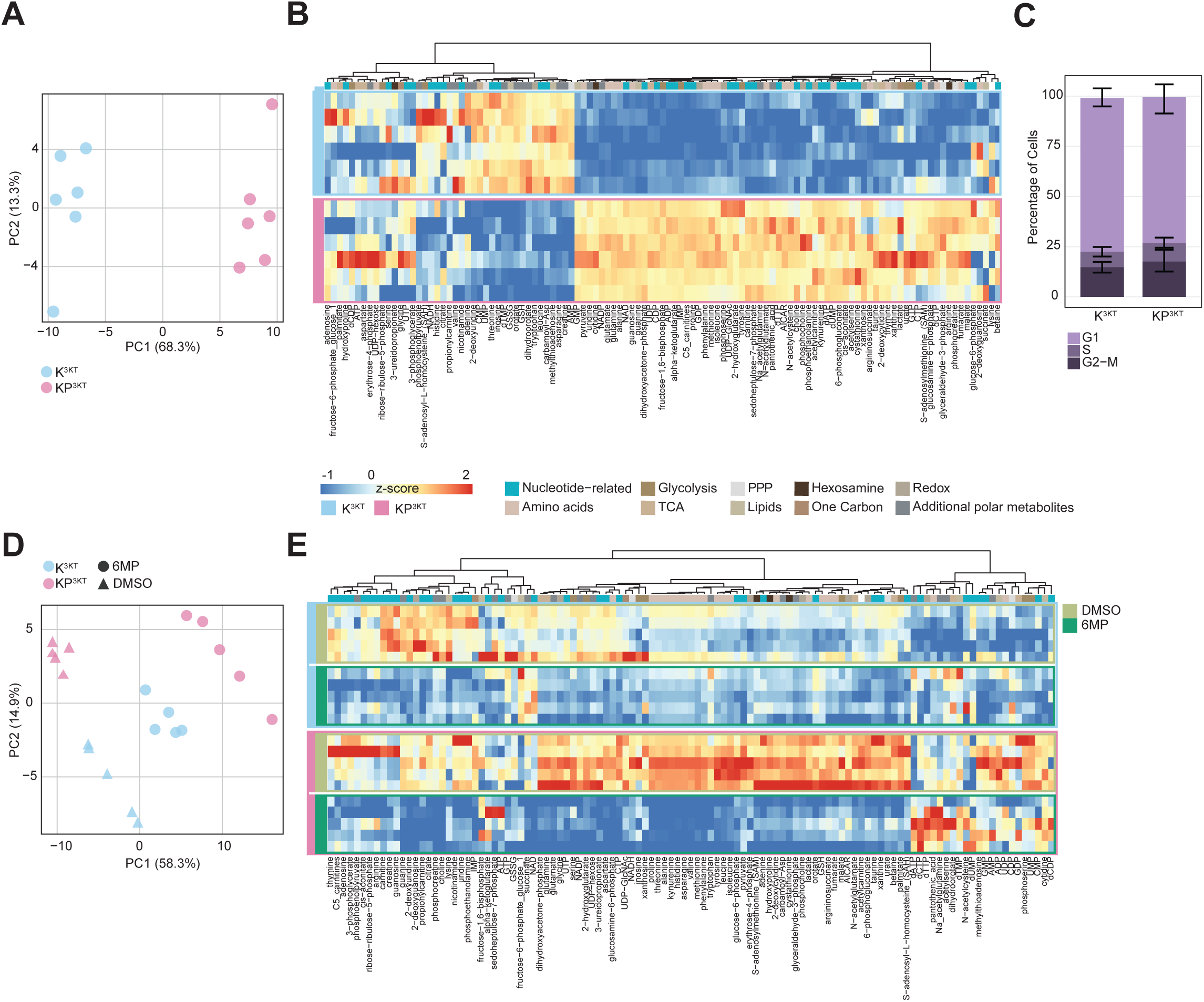
TP53 loss alters purine nucleotide pools without affecting cell cycle progression. A) PCA of Z-scores of polar metabolites in K^3KT^ and KP^3KT^ cells. Pairwise t-test with Bonferroni adjustment. Data are presented as Mean ± SD. P-values: ****P ≤ 0.00005; ***P ≤ 0.0005; **P ≤ 0.005; *P ≤ 0.05. B) Hierarchical clustering heatmap of Z-scores for nucleotide-related and other annotated metabolite classes in K^3KT^ and KP^3KT^ cells. PPP is Pentose Phosphate Pathway. TCA is Tricarboxylic Acid Cycle. Redox is Reduction-Oxidation Pathway. C) Flow cytometry quantification of cell cycle distribution (G1, S, G2-M) in K^3KT^ and KP^3KT^ cells. Pairwise t-test with Bonferroni adjustment. Data are presented as Mean ± SD. P-values: ****P ≤ 0.00005; ***P ≤ 0.0005; **P ≤ 0.005; *P ≤ 0.05. D) PCA of Z-scores of polar metabolites in K^3KT^ and KP^3KT^ cells treated with 5 µM 6MP or DMSO for 72 hours. Pairwise t-test with Bonferroni adjustment. Data are presented as Mean ± SD. P-values: ****P ≤ 0.00005; ***P ≤ 0.0005; **P ≤ 0.005; *P ≤ 0.05. E) Hierarchical clustering heatmap of Z-scores for nucleotide-related and other metabolite classes in K^3KT^ and KP^3KT^ cells following 5 µM 6MP or DMSO treatment for 72 hours. Data represent N=5 (A-C) or N=6 (E-F) independent experiments.

**Supplementary Figure 4.**
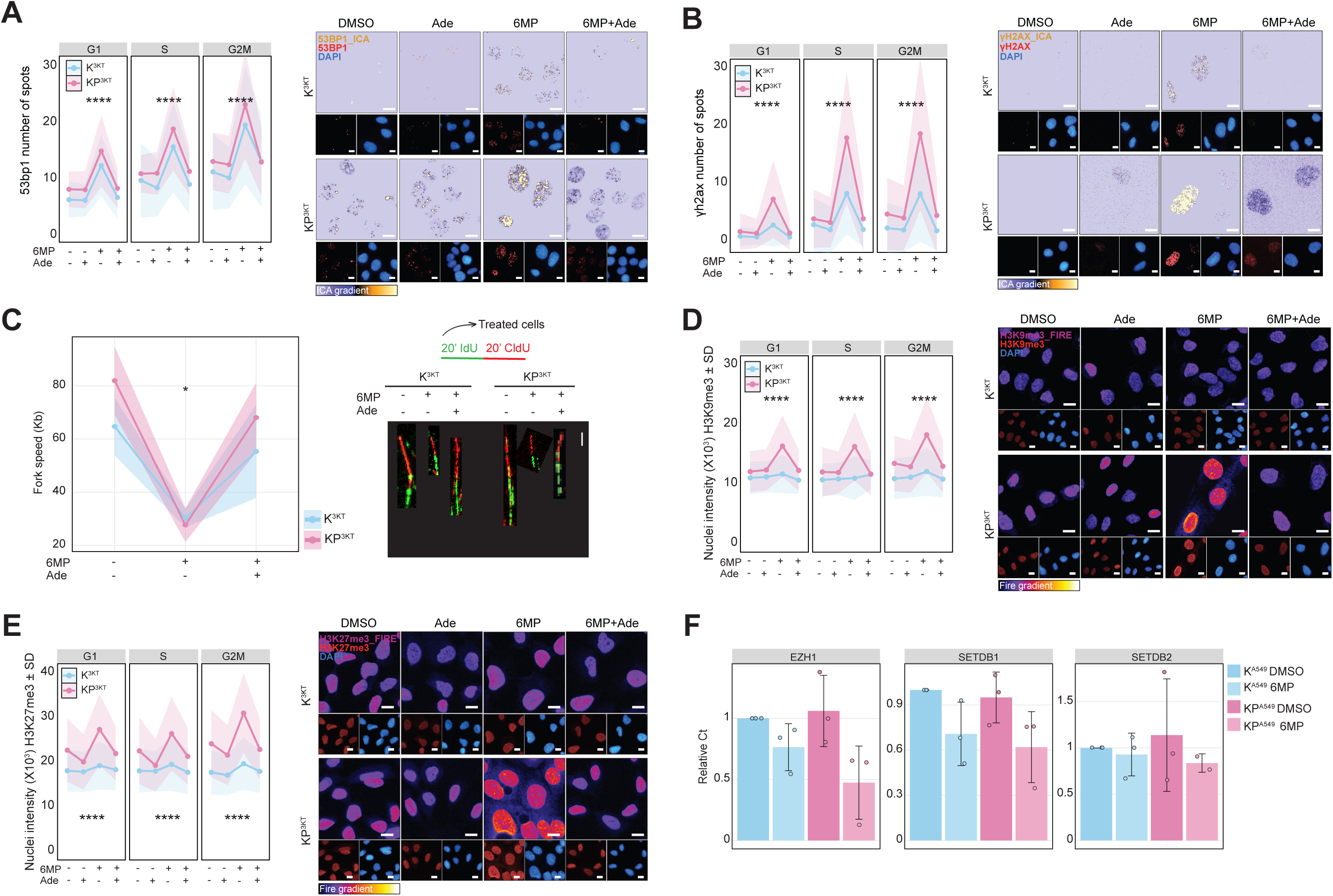
Purine depletion induces DNA damage, impairs replication fork progression, and alters histone methylation, with enhanced sensitivity in TP53-deficient cells. K^3KT^ and KP^3KT^ cells treated with either DMSO or 5 µM 6MP for 72 hours, in presence or absence of 100 µM Adenosine. A) (left) Quantification of 53BP1 foci, (right) representative images. B) (left) Quantification of γH2AX foci, (right) representative images. 53BP1 and γH2AX are in ICA_gradientred and DAPI marks nuclei in blue. Scale bar = 10 µm. C) DNA fibre assay. (left) Quantification of replication fork speed of K^3KT^ and KP^3KT^ cells, (right) representative combed DNA fibres. Mean ± SD, two-tailed unpaired t-test, *P ≤ 0.05. N=20-41 fibres were analysed per condition. Scale bar = 10 µm. D) (left) Quantification of H3K9me3 nuclear intensity and (right) representative images in K^3KT^ and KP^3KT^ cells between different conditions. E) (left) Quantification of H3K27me3 nuclear intensity and (right) representative images. 53BP1 and γH2AX are in ICA_gradientred and DAPI marks nuclei in blue. Scale bar = 10 µm. N=1000-4000 cells. Data are presented as mean ± SD of two independent experiments (N=2). Bonferroni adjusted one-way ANOVA. ****P ≤ 0.00005.

**Supplementary Figure 5.**
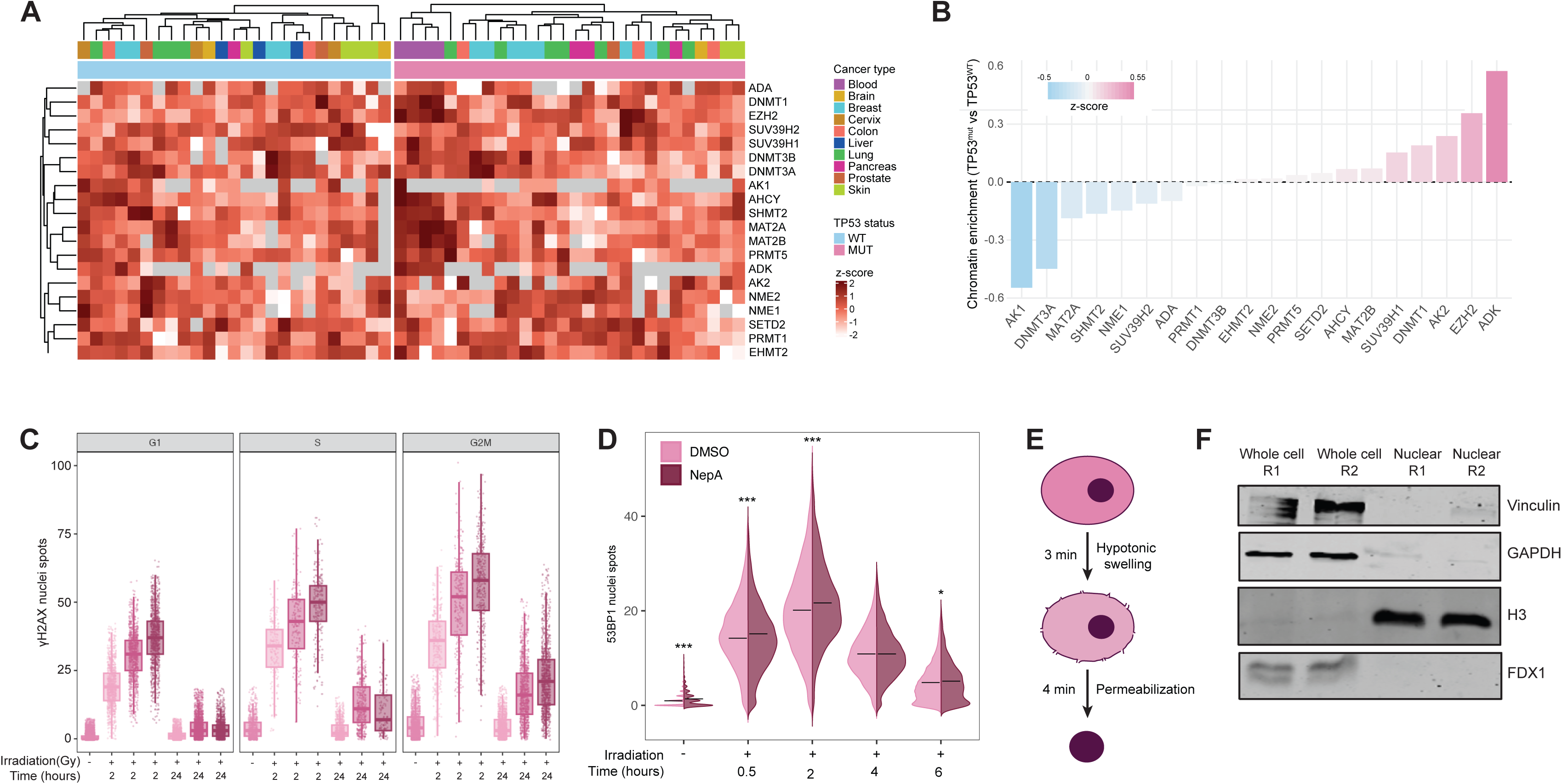
TP53 loss induces DNA methylation to reroute nuclear adenosine recycling. A) Chromatin abundance heatmap of enzymes involved in SAM associated with adenosine and ATP metabolism and B) enrichment of these enzymes in TP53-deficient cells in comparison to TP53-proficient cells. C) Quantification of γH2AX foci in KP^A549^ cells γ-irradiated with 2, 5 and 10 Gy at 2 and 24 hours post-irradiation. D) Quantification of 53BP1 foci of KP^A549^ cells treated with 10 µM NepA or DMSO and combined with or without γ-irradiation at the indicated timepoints. Two-way ANOVA with Holm-adjusted pairwise comparisons, ****P ≤ 0.00005. E) Schematic representation of nuclei isolation protocol. F) Western blot analysis of vinculin, GAPDH, H3 and FDX1of whole cell and nuclei extracts.

**Supplementary Figure 6.**
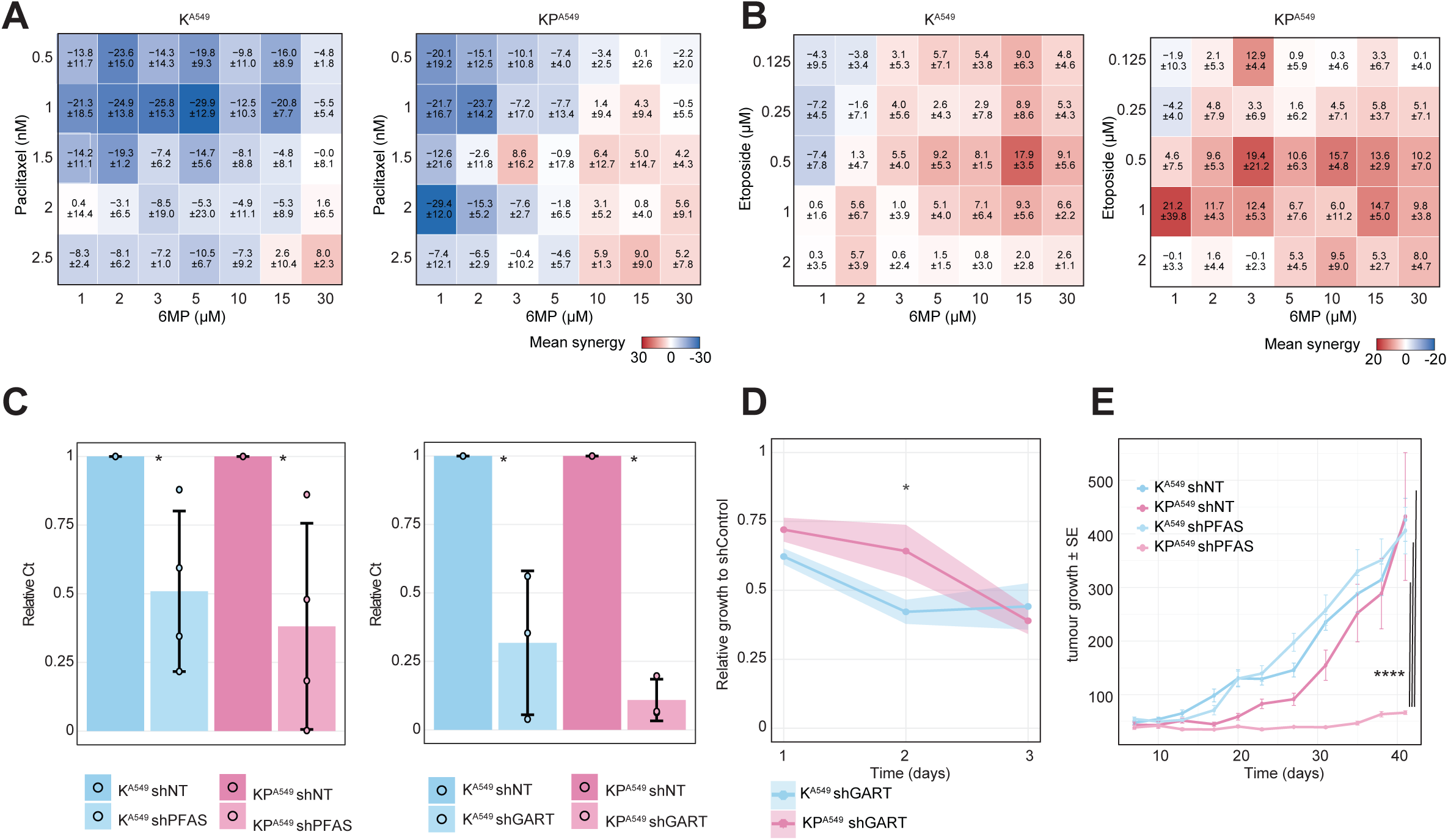
GART depletion similarly affects cell growth in TP53-deficient and TP53-proficient cells. Average Bliss Score ± standard deviation for K^A549^ and KP^A549^ treated with 6MP in combination with A) paclitaxel and B) etoposide with the corresponding concentrations. C) qRT-PCR analysis of PFAS and GART expression in K^A549^ and KP^A549^ cells that transduced with either an shControl, shPFAS or shGART. GAPDH expression was used for data normalisation. Mean ± SD. N=3-4 replicates. Pairwise t-test with Benjamini-Hochberg adjusted. *P ≤ 0.05. D) Batch corrected relative cell growth of K^A549^ and KP^A549^ cells transduced with shGART, normalised to shControl and analysed by HT-IF for nuclei counting. N=3. Bonferroni adjusted Wilcoxon pairwise comparison. *P ≤ 0.05. E) Tumour volume of indicated genotypes was calculated at the experimental endpoint. Data are presented as Mean ± SEM. ****P ≤ 0.00005; p-values were calculated with Linear Mixed Effect Model with POST-HOC Tukey HSD correction.

## Authors’ contributions

J.U.S., A.B.F., Conceptualization. Data curation. Formal analysis. Investigation. Methodology. Validation. Visualization. Project administration. Writing - original draft. Writing - review & editing. A.S.Z. Conceptualization. Data curation. Formal analysis. Investigation. Methodology. Validation. Visualization. Writing - review & editing. L.O., A.C.M., C.M.M, Methodology. Conceptualization. Data curation. Writing - review & editing. S.K., E.A, C.R.E Methodology, Investigation. Writing - review & editing. M.G.V.H. Conceptualization. Visualization. Writing - original draft. Writing - review & editing. S.S. and A.J. Conceptualization. Visualization. Project administration. Writing - original draft. Writing - review & editing. Supervision. Resources. Funding acquisition.

## Acknowledgements

We thank Dr Lluis Espinosa, Dr Ivano Amelio, Dr S. Aranda, Dr. H. Imamura, Dr A.F. Meyerhans, Dr. J.M. Díez Antón and Dr S. Vicent Cabra for providing reagents; and Dr K. Sutherland and Dr J. Valcarcel for discussions; CRG/UPF Flow Cytometry Unit and Raul Pena with sample preparation. This work was supported by grants and fellowships from the Spanish Ministry of Science and Innovation Grant to A.J. (PID2021-127710OB-I00, CNS2023-144819), “La Caixa” foundation to A.J. (51110009 and HR22-00402). S.S. acknowledges financial support to this project from the European Research Council (ERC) under the European Union’s Horizon 2020 research and innovation programme (grant agreement no. 852343). J.U.S acknowledges financial support from the “La Caixa” foundation Doctoral INPhINIT Retaining Fellowships (LCF/BQ/DR22/11950019). A.B.F. acknowledges financial support from the AGAUR-FI doctoral Fellowship 2025 STEP 00304. A.S.Z acknowledges financial support from the Longitude Capital fellowship (T32GM007287). A.J. is supported by Ramon y Cajal Research Fellowship (RYC2018-025244-I). This work was made possible through the “Unidad de Excelencia María de Maeztu’’ funded by the MCIN and AEI (CEX2018-000792-M). M.G.V.H. acknowledges funding from the NCI R35CA242379 and P30CA014051 as well as the Ludwig Center at MIT, Breakthrough Cancer, and the MIT Center for Precision Cancer Medicine.

